# An Issue of Concern: Unique Truncated ORF8 Protein Variants of SARS-CoV-2

**DOI:** 10.1101/2021.05.25.445557

**Authors:** Sk. Sarif Hassan, Vaishnavi Kodakandla, Elrashdy M. Redwan, Kenneth Lundstrom, Pabitra Pal Choudhury, Tarek Mohamed Abd El-Aziz, Kazuo Takayama, Ramesh Kandimalla, Amos Lal, Ángel Serrano-Aroca, Gajendra Kumar Azad, Alaa A. A. Aljabali, Giorgio Palu, Gaurav Chauhan, Parise Adadi, Murtaza Tambuwala, Adam M. Brufsky, Wagner Baetas-da-Cruz, Debmalya Barh, Nicolas G Bazan, Vladimir N. Uversky

## Abstract

Open reading frame 8 (ORF8) protein is one of the most evolving accessory proteins in severe acute respiratory syndrome coronavirus 2 (SARS-CoV-2), the causative agent of coronavirus disease 2019 (COVID-19). It was previously reported that the ORF8 protein inhibits presentation of viral antigens by the major histocompatibility complex class I (MHC-I) and interacts with host factors involved in pulmonary inflammation. The ORF8 protein assists SARS-CoV-2 to evade immunity and replication. Among many contributing mutations, Q27STOP, a mutation in the ORF8 protein defines the B.1.1.7 lineage of SARS-CoV-2, which is engendering the second wave of COVID-19. In the present study, 47 unique truncated ORF8 proteins (T-ORF8) due to the Q27STOP mutations were identified among 49055 available B.1.1.7 SARS-CoV-2 sequences. The results show that only one of the 47 T-ORF8 variants spread to over 57 geo-locations in North America, and other continents which includes Africa, Asia, Europe and South America. Based on various quantitative features such as amino acid homology, polar/non-polar sequence homology, Shannon entropy conservation, and other physicochemical properties of all specific 47 T-ORF8 protein variants, a collection of nine possible T-ORF8 unique variants were defined. The question of whether T-ORF8 variants work similarly to ORF8 has yet to be investigated. A positive response to the question could exacerbate future COVID-19 waves, necessitating severe containment measures.

## 1. Introduction

The world is proceeding through an unprecedented time due to the Coronavirus disease 2019 (COVID-19), of which the causative agent is the severe acute respiratory syndrome coronavirus 2 (SARS-CoV-2) [1, 2, 3, 4, 5]. There are nine open reading frames (ORFs), which encodes for accessory proteins important for the modulation of the metabolism in infected host cells and innate immunity evasion via a complicated signalome and an interactome [6, 7, 8, 9, 10]. The ORF8 protein is one of the most rapidly evolving accessory proteins among the beta coronaviruses, not only due to its ability to interfere with host immune responses [11, 12, 13, 14]. It directly interacts with major histocompatibility complex class I (MHC-I) both invitro and invivo, and is down-regulated, which impairs its ability to antigen presentation and rendering infected cells less sensitive to lysis by cytotoxic T lymphocytes [15]. ORF8 suppresses type I interferon antiviral responses and interacts with host factors involved in pulmonary inflammation and fibrogenesis [15, 16]. From all viral proteomes interacting with human metalloproteome, the ORF8 interplay with 10 out 58 [17]. ORF8 (of SARS-CoV-2 and SARS-CoV) play crucial roles in virus pathophysiological events, it dysregulates the TGF-β pathway, which is involved in tissue fibrosis [18]. The functional implications of SARS-CoV-2 ORF8 had already gained huge attention and ORF8 is considered an important component of the immune evasion machinery [11, 18, 19, 20]. The SARS-CoV-2 ORF8 protein has less than twenty percent amino acid sequence homology with the SARS-CoV ORF8, and is a rapidly evolving protein [14, 21]. A molecular framework for understanding the rapid evolution of ORF8, its contributions to COVID-19 pathogenesis, and the potential for its neutralization by antibodies were supported by the structural analysis of the ORF8 protein [22, 23]. The crosstalk between viral (SARS-CoV-2 or SARS-CoV) infections and host cell proteome at different levels may enable identification of distinct and common molecular mechanisms [15]. Of note, SARS-CoV-2 ORF8 ORF8 not only interacts with a significant number of host proteome related to endoplasmic reticulum quality control, glycosylation, and extracellular matrix organization, although the mechanism of action of ORF8 concerning those interacting proteins is uncertain, so far [23, 24].

The clade S, a subtype of SARS-CoV-2, was identified to possess the mutation L84S in the ORF8 protein sequence [25, 26, 27]. Presently, among many variants of SARS-CoV-2, the lineage B.1.1.7 carries a larger than usual number of genetic changes [28, 29, 30]. Among many non-synonymous mutations, Q27STOP in the ORF8 protein contributed to deduce the branch leading to lineage B.1.1.7 [31, 32]. The Q27STOP mutation inactivates ORF8 protein favoring further downstream mutations and could be responsible for the increased transmissibility of the B.1.1.7 variant [28, 33]. The B.1.1.7 variant was found to be more transmissible than the wild-type SARS-CoV-2 and was first detected in September 2020 in the UK [34, 35]. Further, it began to spread rapidly by mid-December, and is correlated with a significant increase in SARS-CoV-2 infections in the UK and worldwide.

Functional implications on the immune surveillance of ORF8 due to the truncation at position 27 remain unclear [18]. Thus, it is of utmost importance to gain insight into the functionality of the truncated ORF8 protein variants to comprehend the B.1.1.7 lineage through theoretical and experimental characterization and genomic surveillance worldwide [36]. The present study was aimed to characterize the unique variations of truncated ORF8 proteins (T-ORF8) due to the Q27STOP mutation. Further, this investigation differentiates a single T-ORF8 variant among 47 distinct unique T-ORF8 protein variants present in SARS-CoV-2, worldwide as of May 20th, 2021. Several clusters of the unique T-ORF8 have been identified based on various bioinformatics features and phylogenetic relationships, along with emerging variants of the unique T-ORF8.

## 2. Data acquisition and methods

Truncated ORF8 protein (T-ORF8) sequences (complete) from five continents (Asia, Africa, Europe, South America, and North America) were downloaded in Fasta format (as of May 18, 2021) from the National Center for Biotechnology Information (NCBI) database (http://www.ncbi.nlm.nih.gov/). Note that no T-ORF8 protein sequence was found from Oceania as of May 18th, 2021. Further, Fasta files were processed in *Matlab-2021a* for extracting unique T-ORF8 sequences for each continent.

### 2.1. Derivation of polar/non-polar sequences and associated phylogeny

Every amino acid in a given T-ORF8 sequence was identified as polar (Q) and non-polar (P). Thus, every unique T-ORF8 became a binary sequence with two symbols P and Q. Then sequence homology of these sequences was derived using the Clustal Omega web-suite and then associated with nearest neighborhood phylogenetic relationship among the unique T-ORF8 variants. Further, unique T-ORF8 variants having distinct binary polar/non-polar sequences were extracted [37, 38].

### 2.2. Frequency distribution of amino acids and phylogeny

The frequency of each amino acid present in a T-ORF8 sequence was determined using standard bioinformatics routine in *Matlab-2021a.* For each T-ORF8 protein, a twenty-dimensional frequency-vector considering the frequency of standard twenty amino acids can be obtained. Based on this frequency distribution of amino acids several consequences were drawn. The distance (Euclidean metric) between any two pairs of frequency vectors was calculated for each pair of T-ORF8 sequences. By having the distance matrix, a phylogenetic relationship was developed based on the nearest neighbor-joining method using the standard routine in *Matlab-2021a* [39, 40].

### 2.3. Amino acid conservation Shannon entropy

The degree of conservation of amino acids embedded in a T-ORF8 protein was obtained by the well-known information-theoretic measure called ‘*Shannon entropy*(SE)’. For each T-ORF8 protein, Shannon entropy of amino acid conservation over the amino acid sequence of T-ORF8 protein was calculated using the following formula [39, 41]:

For a given T-ORF8 sequence of length *l* (here *l* = 26), the conservation of amino acids was calculated as follows:

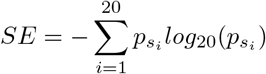

where 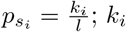; *k_i_* represents the number of occurrences of an amino acid *s_i_* in the T-ORF8 sequence [42].

### 2.4. Prediction of molecular and physicochemical properties

Theoretical pI (PI), extinction coefficient (EC), instability index (II), aliphatic index (AI), protein solubility (PS), grand average of hydropathicity (GRAVY), and the number of tiny, small, aliphatic, aromatic, non-polar, polar, charged, basic and acidic residues of all unique T-ORF8 proteins were calculated using the web-servers ‘ProtParam’, ‘Protein-sol’ and EMBOSS Pepstats [43, 44, 45].

### 2.5. Intrinsic disorder analysis

All 47 T-ORF8 variants were subjected to the per-residue disorder analysis, for which PONDR-VSL2 algorithm was employed [46]. This tool shows good performance on proteins containing both structure and disorder and was favorably ranked in a recent Critical Assessment of protein Intrinsic Disorder prediction (CAID) experiment [47].

### 2.6. Finding functional motifs

The Eukaryotic Linear Motif (ELM) resource (http://elm.eu.org/) was used for finding functional sites in proteins [48]. ELMs (also known as short linear motifs (SLiMs)), are short protein interaction sites, which are commonly found in intrinsically disordered regions of proteins and define a wide range of protein functionality.

## 3. Results

Continent-wise, all unique T-ORF8 protein variants were segregated from a set of available truncated ORF8 protein sequences collected from the NCBI database. Further, variability and commonality of the unique T-ORF8 proteins were analyzed from various quantitative measures such as amino acid homology-based phylogeny, frequency distribution of amino acids and associated phylogeny, polarity sequence-based phylogeny, and physicochemical properties. Relying on these features, a set of nine possible unique T-ORF8 variants were identified, which were found to lie within the likelihood of a T-ORF8 variant named P15 (Table 3).

### 3.1. Characteristics of the unique variants of T-ORF8

For each continent, the number of total sequences, the unique truncated ORF8 (T-ORF8) sequences and percentages are presented in Table 1.

**Table 1:** Frequency and percentages unique T-ORF8 variants (continent-wise)

| *Percentages of the unique T-ORF8 variants on continents* |  |  |  |  |
| --- | --- | --- | --- | --- |
| Continent | *Total T-ORF8 (T)* | *Unique T-ORF8 (U)* | *Percentage, continent-wise* | *Percentage, worldwide* |
| **Africa** | 108 | 1 | 0.926 | 1.96 |
| **Asia** | 99 | 1 | 1.01 | 1.96 |
| **Europe** | 156 | 1 | 0.641 | 1.96 |
| **South America** | 1 | 1 | 100 | 1.96 |
| **North America** | 48691 | 47 | 0.096 | 92.16 |
| **Worldwide** | 49055 | 47 | 0.104 |  |

The results showed that 47 unique T-ORF8 proteins were present in North America. The unique T-ORF8 variants from Africa, Asia, Europe, and South America were contained in the set of unique T-ORF8 variants available in North America.

Additionally, there were seven T-ORF8 with amino acid lengths 22, 24, 40 and 41 as of May 18, 2021 available in North America (Table 2).

**Table 2:**
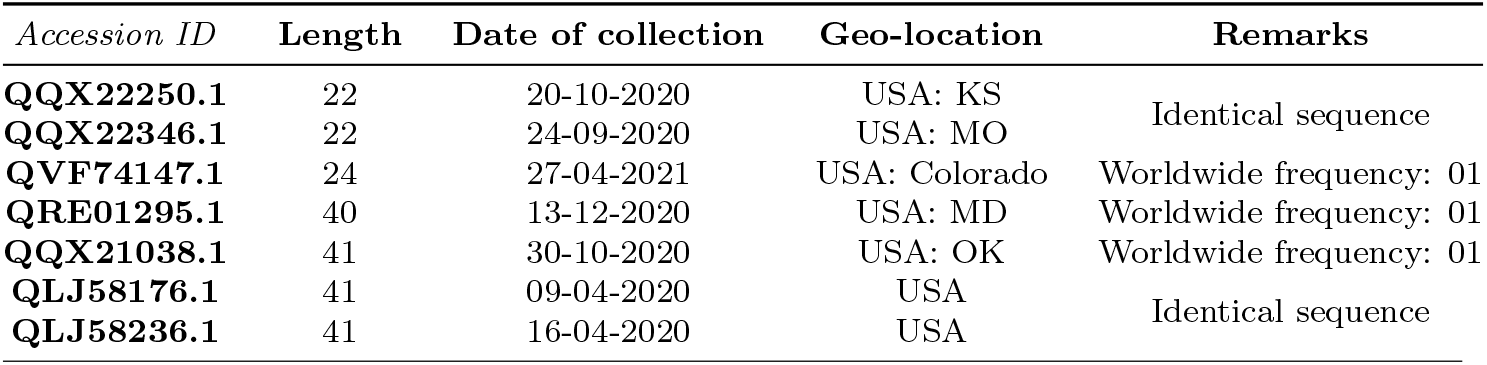
Truncated ORF8 variants of length other than 26

Note that among the seven T-ORF8 sequences, only five were found to be unique as mentioned in Table 2. As of May 18, 2021 a single copy of the T-ORF8 proteins of amino acid lengths of 24 and 41 (Table 2) were found. There were two T-ORF8 variants of 41 amino acids available in North America. The most frequent T-ORF8 proteins so far observed were the T-ORF8 proteins of 26 amino acids. It was observed that the T-ORF8 arose due to truncation at the residue positions 23, 25, 27, 41, and 42 of the complete ORF8 protein (121 aa long sequence). We investigated the possible mutations for such truncations. A snapshot of the amino acid residues and their possible mutations with respect to the reference sequence NC_045512 is presented in Figure 1.

**Figure 1:**
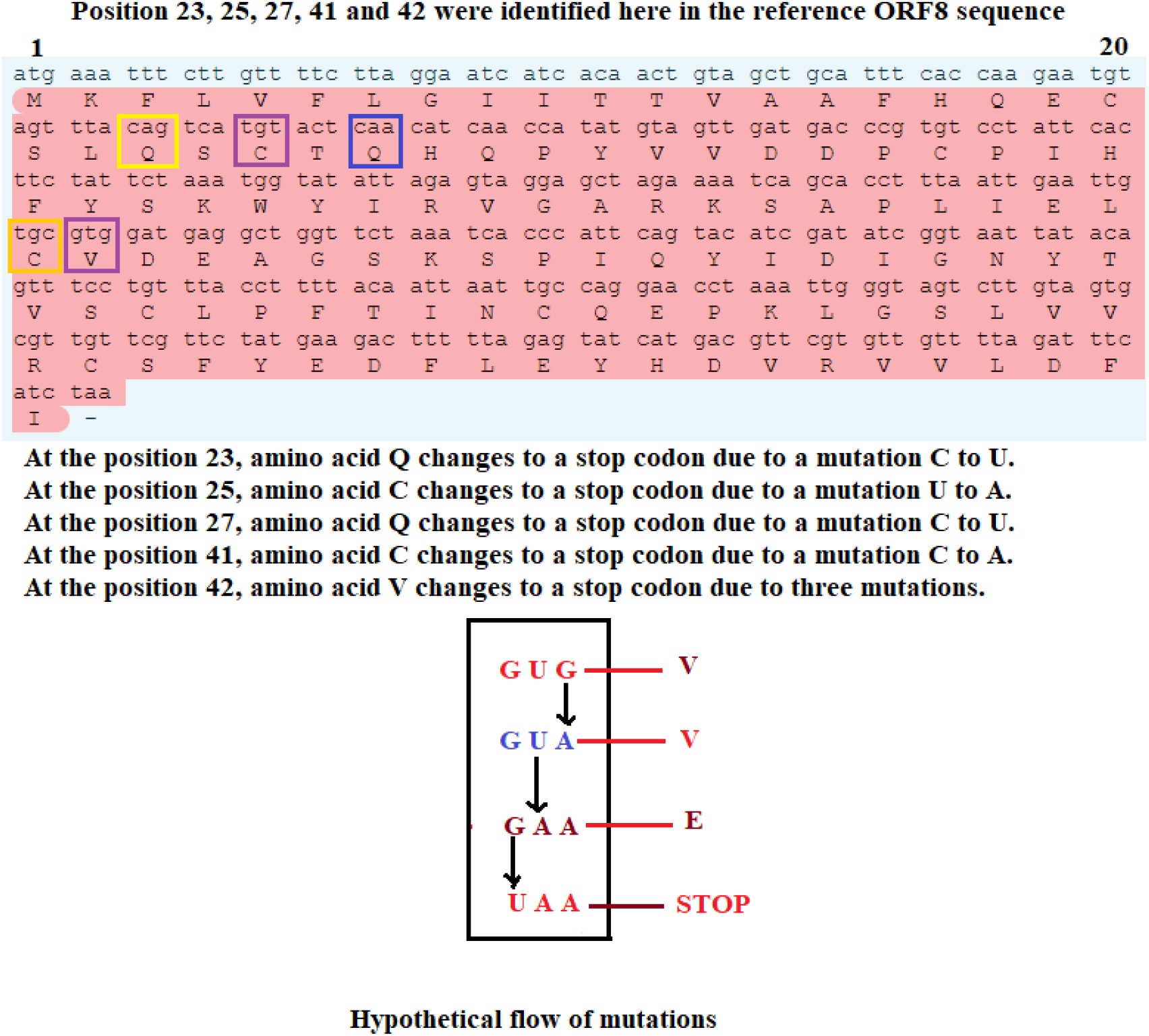
Possible mutations for truncation at 23, 25, 27, 40, and 42 residue position of ORF8 protein (NC_045512) of SARS-CoV-2.

Note that in four positions 23, 25, 27 and 40 amino acids Q and C both were truncated due to mutations at the first and third position, respectively, of the respective codon. The amino acid Valine (V) was truncated due to three mutations at the third, second and first positions of the codon ‘GUG’. Furthermore, it was observed that the mutations at the positions 23 and 25 were identical (C to U) and the changes of bases were transition mutations i.e., pyrimidine (purine) to pyrimidine (purine), whereas the changes of bases of the truncated mutations at positions 25 and 41 were transversal mutations i.e. pyrimidine (purine) to purine (pyrimidine). For position 42, three sequences of mutations were hypothesized, taking place at first, second, and third positions of the codon (GUG) i.e., transition mutations (purine to purine), transversal mutation (pyrimidine to purine), and transversal mutation (purine to pyrimidine) respectively.

The list of unique T-ORF8 sequences of 26 amino acids with their representative accession IDs and sequences is presented in Table 3.

**Table 3:**
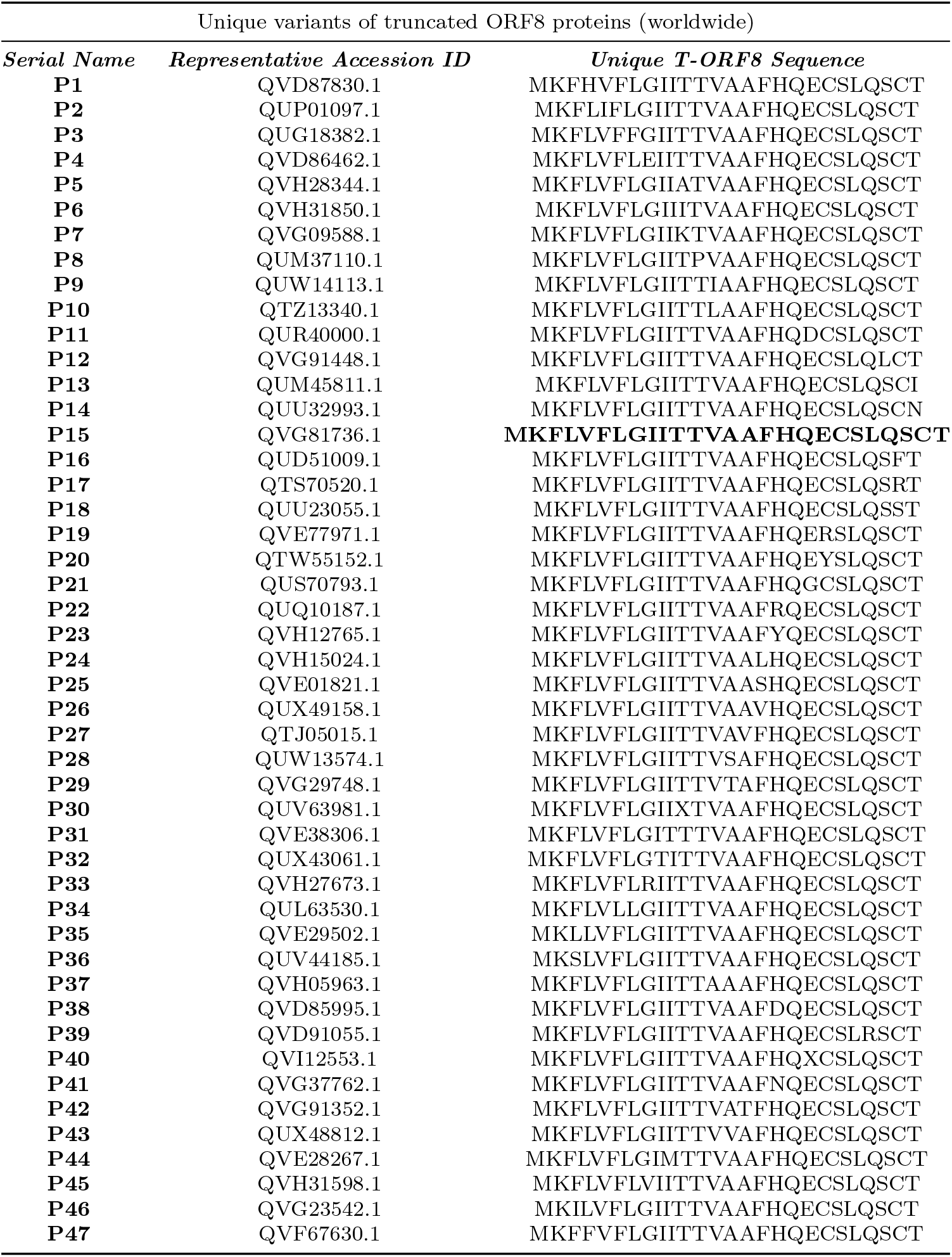
List of unique truncated ORF8 proteins and their representative accession IDs

Further, it was found that the unique T-ORF8 variants from Africa, Asia, Europe and South America were identical with relation to P15, as illustrated in Table 3.

The date of sample collection, geo-location and accession ID of the first identified SARS-CoV-2 containing unique T-ORF8 variants are presented in Table 4.

**Table 4:**
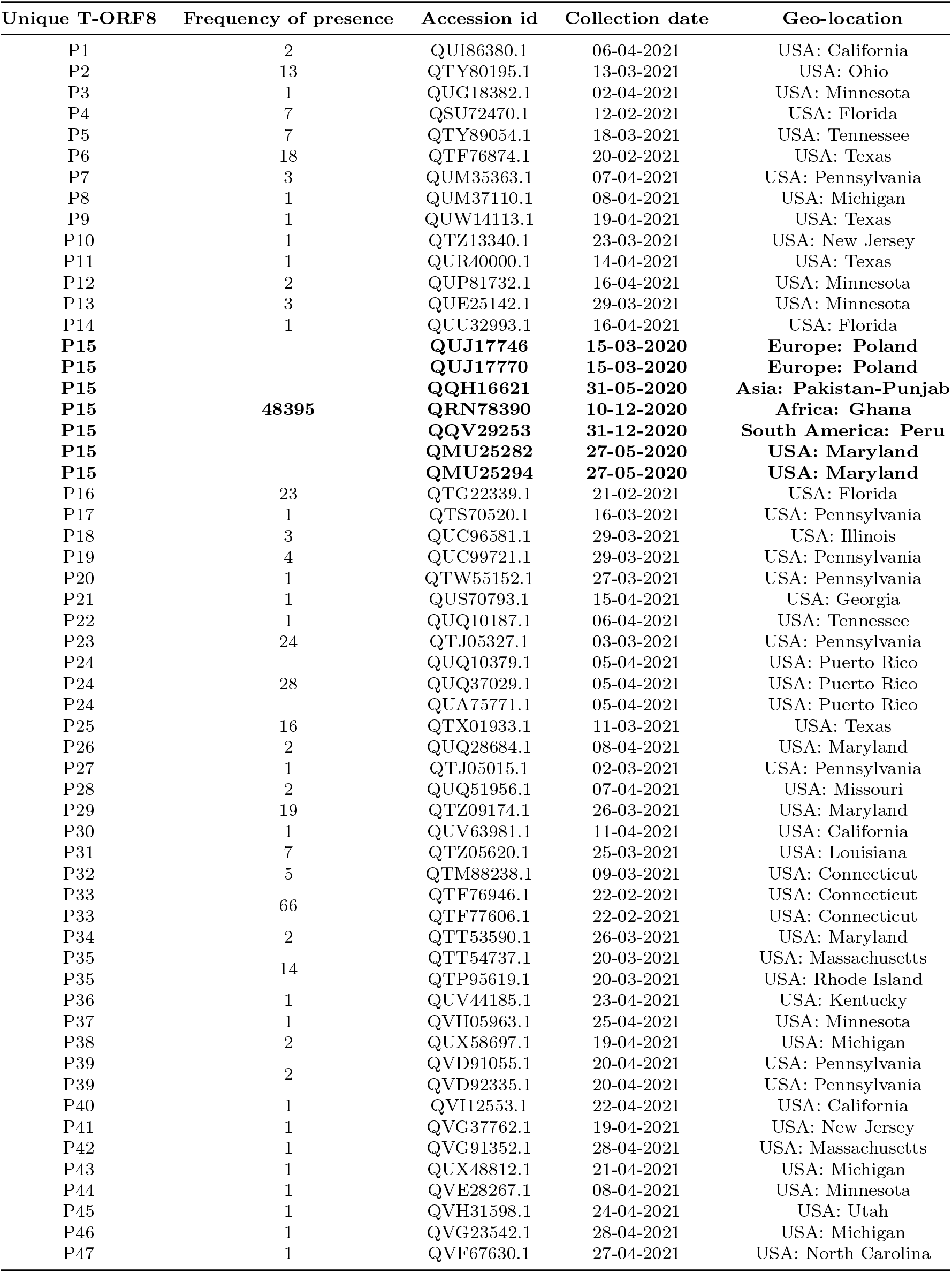
Collection date, geo-location, frequency of presence and accession ID of the first identified of each unique T-ORF8 variants

The ORF8 protein sequence P15 was found in 48395 copies of the B.1.17 SARS-CoV-2 lineage in North America. Besides, the P15 variant having the Q27STOP mutation in the B.1.1.7 lineage was found on Africa, Asia, Europe and South America with frequency 108, 99, 156, and 1 respectively. None of the other 46 T-ORF8 unique variants was found in any continent, as of May 18, 2021. So, 46 unique T-ORF8 sequences were exclusively found in North America. Therefore, the P15 TORF8 variant is of particular interest for its uniqueness due to its apparent prevalence in most of the B.1.17 lineages of SARS-CoV-2 from North America and other continents.

In Europe, the P15 variant was first detected in two infected patients from Poland on March 15, 2020. In North America, on May 27, 2020, two patients from Maryland were infected with the same SARS-CoV-2 P15 variant. After five days of the second occurrence of P15 in North America, one patient from Punjab-Pakistan (Asia) was infected by the P15 SARS-CoV-2 variant. Six months thereafter the same variant was found in a patient from Ghana, for the first time in Africa. Twenty days after the fifth occurrence in Africa, on December 31, 2020, P15 variant was identified in Peru (South America).

Additionally, the frequency distribution of the T-ORF8 P15 variants across the North American continent is presented in Table 5. It was found that the T-ORF8 P15 variant spread over three geo-locations Michigan, Florida and Minnesota with the highest number of frequencies of 5084, 6884, and 7416, respectively.

**Table 5:**
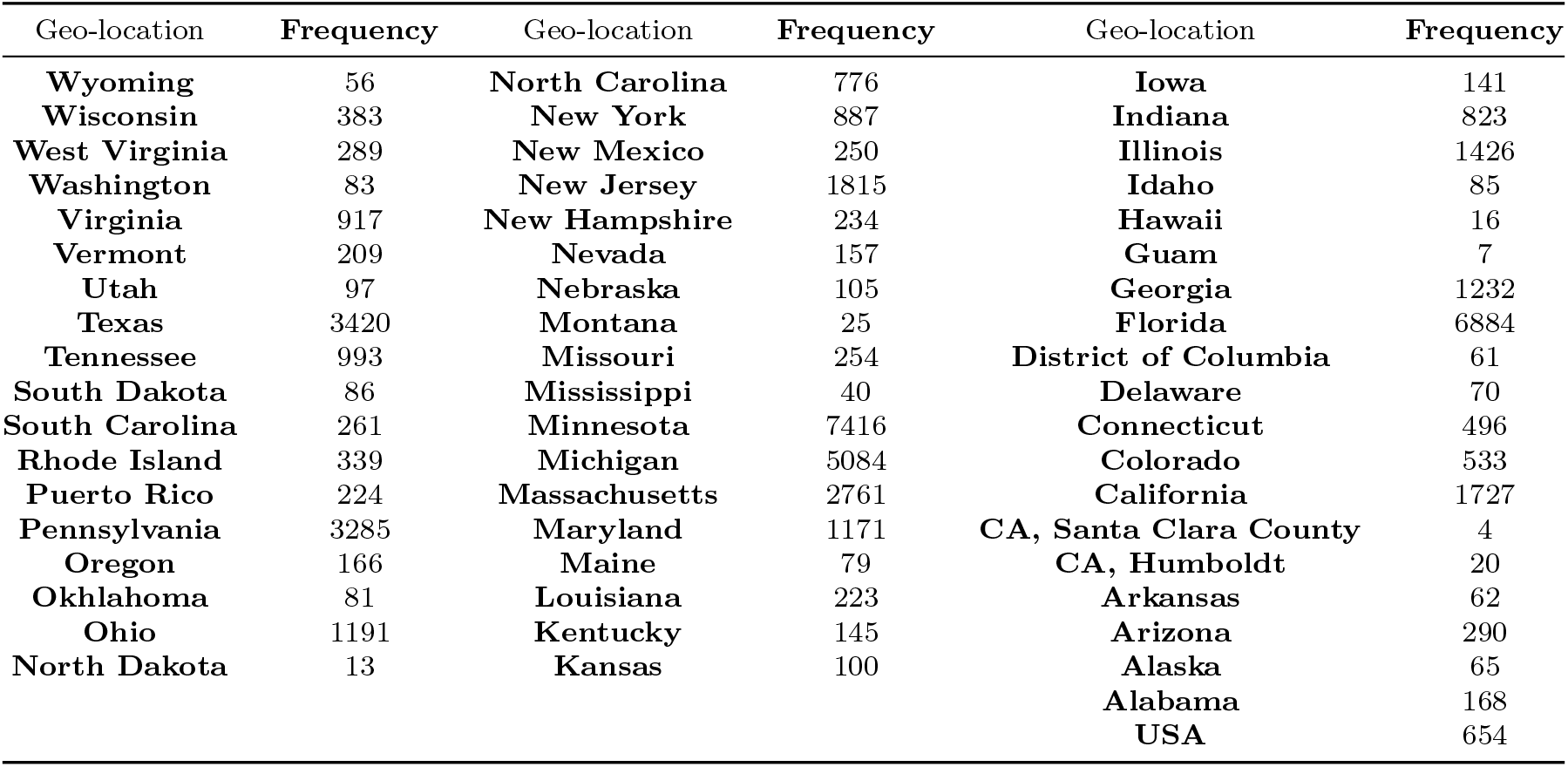
Distribution of cumulative frequency of P15 variants across North America

The P15 variant was found for the first time in Maryland, but the frequency at this geo-location was 1171 on May 14, 2021. There were 18 geo-locations, where the frequency of spread of the P15 variant was found to be less than 100 (Table 5). The frequency distribution of the presence of the T-ORF8 P15 variant in different geo-locations of North America is presented in Figure 2.

**Figure 2:**
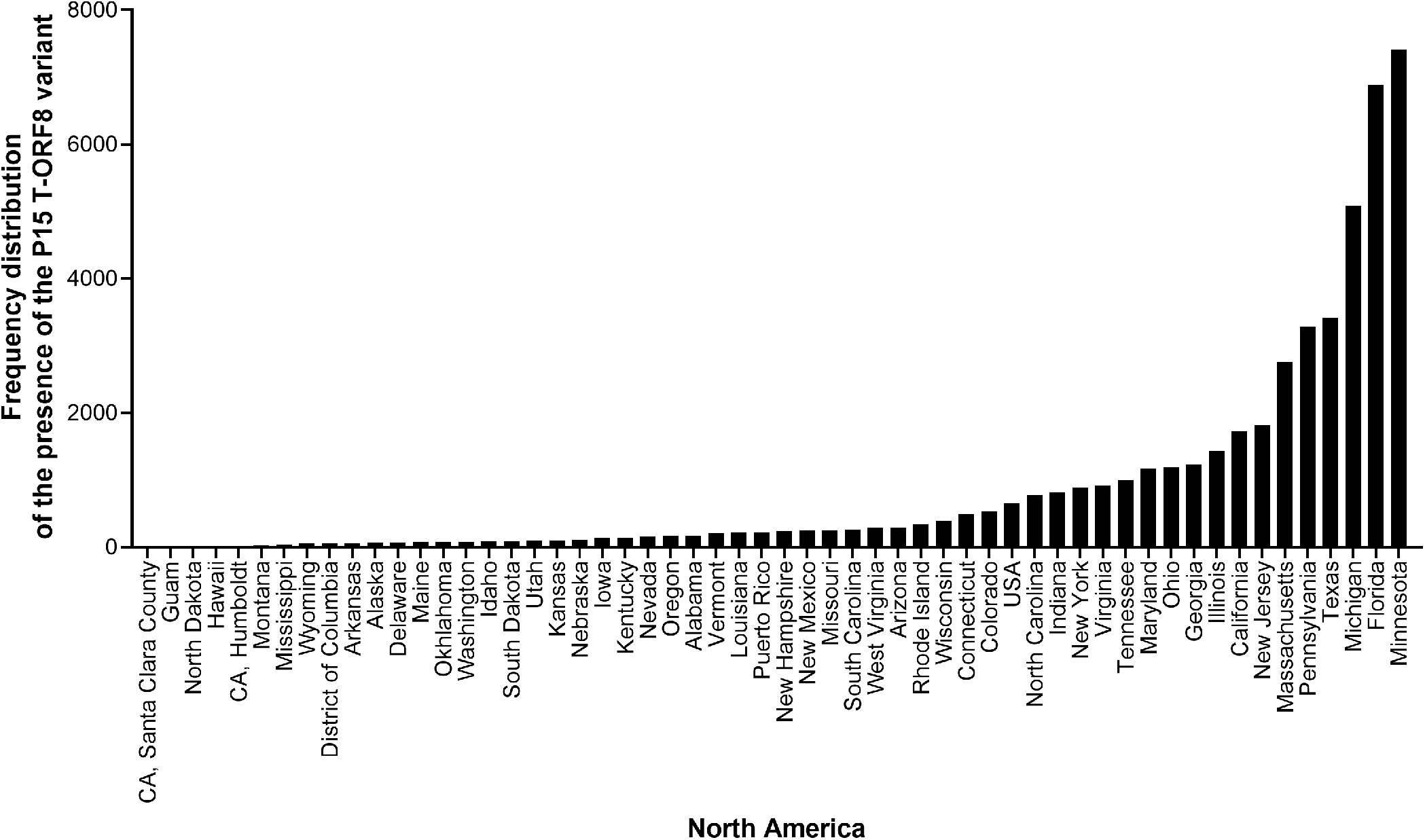
Frequency distribution of the presence of the P15 T-ORF8 variant in different geo-locations across North America.

Among other geo-locations, Guam (US territory located in the Pacific Ocean) and North Dakota, USA had the least number of patients infected by the B.1.1.7 variant containing the P15 protein. In Guam, according to the NCBI SARS-CoV-2 database, all seven patients were infected by the B.1.1.7 variant of SARS-CoV-2 containing the P15, within a short period from February 21 to April 11, 2021. Also, in North Dakota, 13 of 16 patients were infected by the same strain of SARS-CoV-2 from February 2, 2021 to April 28, 2021.

The frequency distribution of all T-ORF8 variants across the US is presented in Table 6. It is evident that all 47 unique T-ORF8 variants were detected in 21 different states of the US.

**Table 6:** Frequency distribution of unique T-ORF8 variants over the USA

| Table 6: Frequency distribution of unique T-ORF8 variants over the USA |  |  |  |
| --- | --- | --- | --- |
| USA: states | Unique T-ORF8 variants | USA: states | Unique T-ORF8 variants |
| USA: California | P1, P30, P40 | USA: Missouri | P28 |
| USA: Connecticut | P32, P33 | USA: New Jersey | P10, P41 |
| USA: Florida | P4, P14, P16 | USA: Ohio | P2 |
| USA: Georgia | P21 | USA: North Carolina | P47 |
| USA: Illinois | P18 | USA: Pennsylvania | P7, P17, P19, P20, P23, P27, P39 |
| USA: Kentucky | P36 | USA: Puerto Rico | P24 |
| USA: Louisiana | P31 | USA: Tennessee | P5, P22 |
| USA: Maryland | P15, P26, P29, P34 | USA: Rhode Island | P35 |
| USA: Massachusetts | P35, P42 | USA: Texas | P6, P9, P11, P25 |
| USA: Michigan | P8, P38, P43, P46 | USA: Utah | P45 |
| USA: Minnesota | P3, P12, P13, P37, P44 |  |  |

The frequency distribution of the unique T-ORF8 variants in 21 states of the US is presented in Figure 3.

**Figure 3:**
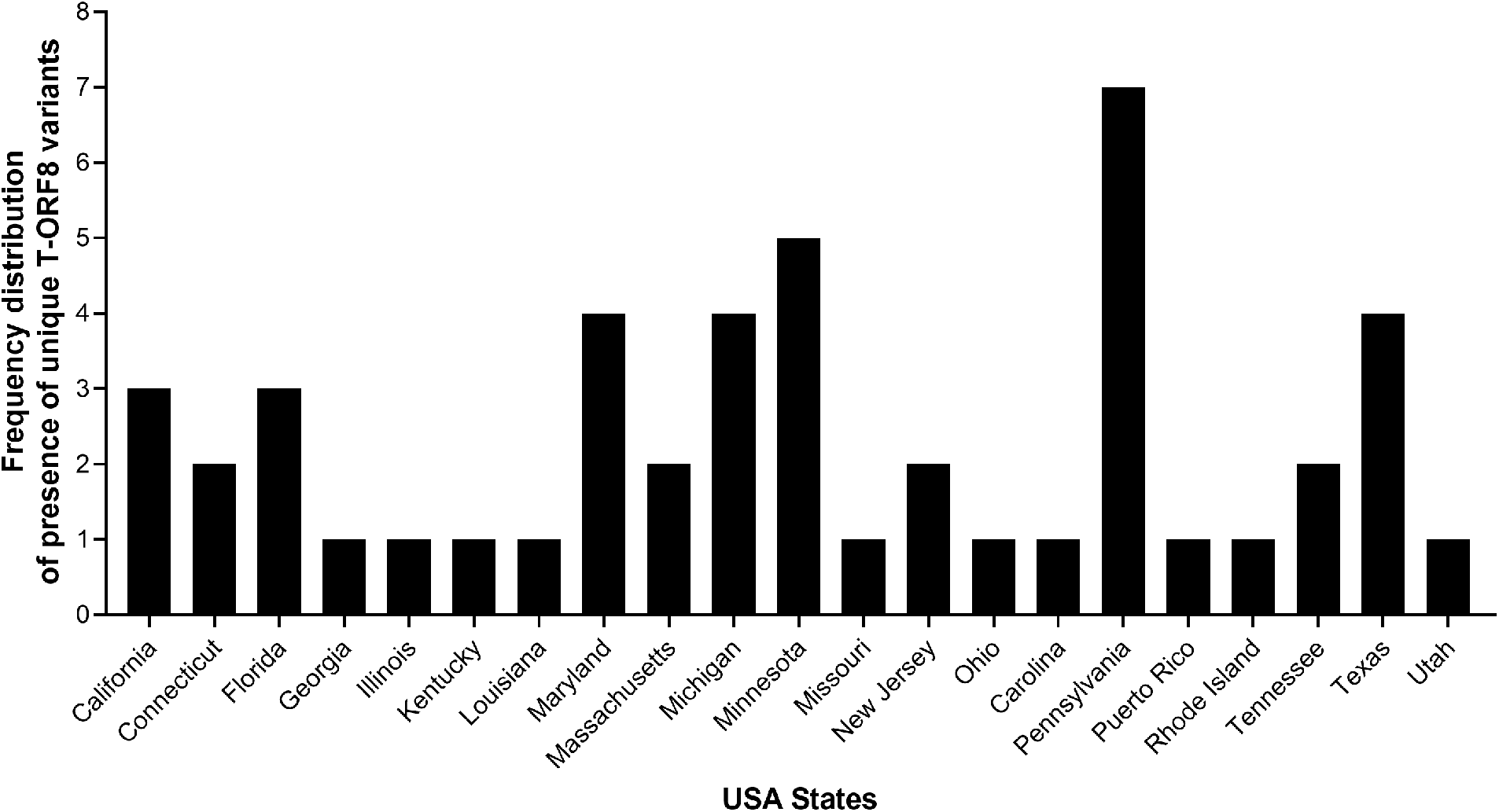
Frequency distribution of the unique T-ORF8 variants in different states of the US.

The highest number (7) of unique T-ORF8 variants was detected as a first instance in Pennsylvania within a short period (March 2 to April 20, 2021). The P35 variant was found initially in two states: Rhode Island and Massachusetts on March 20, 2021. Furthermore, it was observed that all T-ORF8 variants other than P15 emerged for the first time in SARS-CoV-2 from February 12, 2021 to April 28, 2021.

Application the Clustal Omega web-server, an amino acid sequence-based alignment and corresponding phylogenetic tree of the unique T-ORF8 variants are presented in Figure 4.

**Figure 4:**
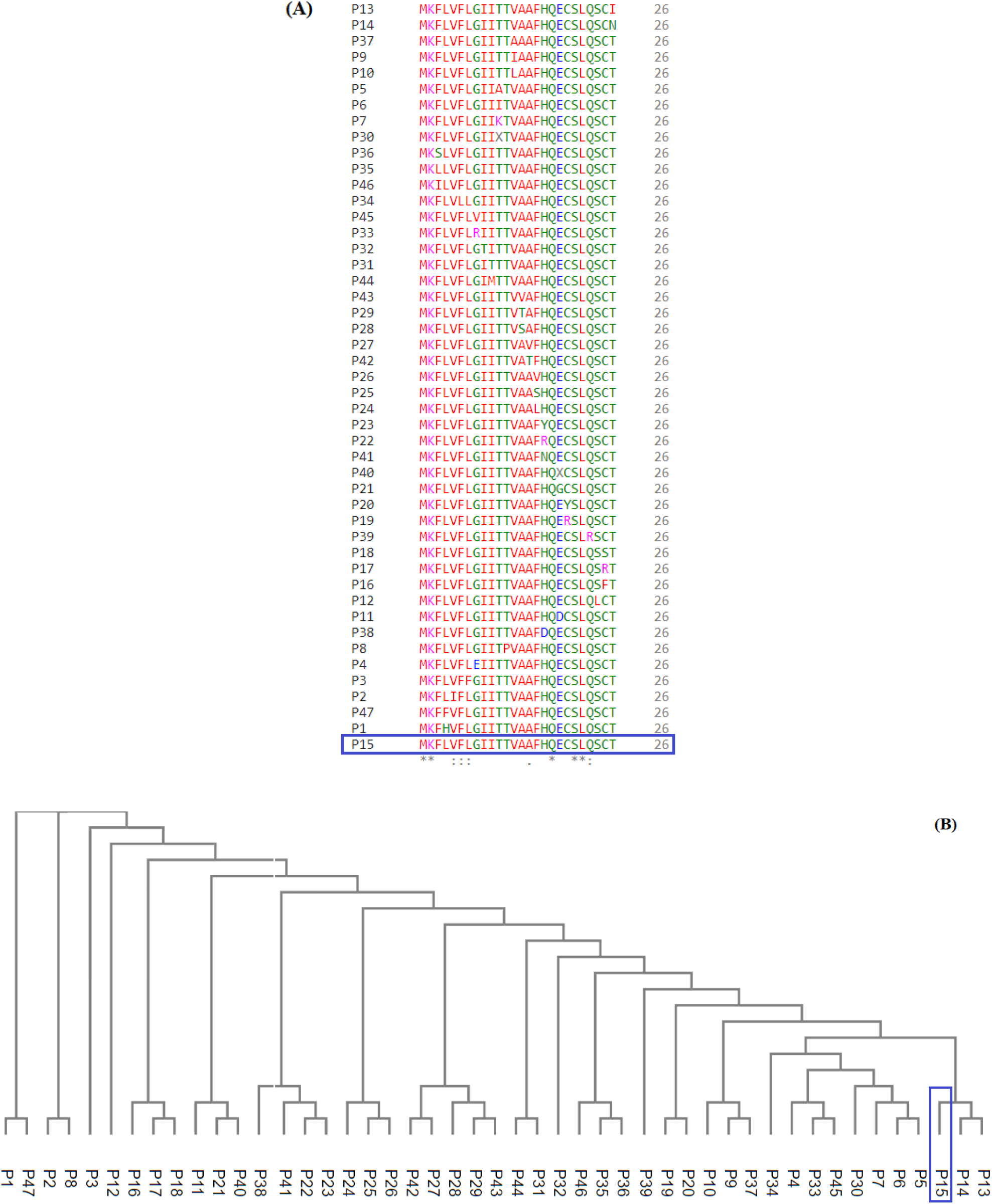
Analysis of variability among unique T-ORF8 variants: (A) Amino acid sequence-based alignment of unique T-ORF8 proteins using Clustal-Omega, and (B) associated phylogenetic relationship among the unique T-ORF8 proteins.

From the sequence alignment it was derived that all unique T-ORF8 variants share identical amino acids M, K, Q, S, and L at the positions 1, 2, 18, 21, 22 respectively. Further it was found that T-ORF8 P15 is much closer to the ORF8 sequences P13 and P14. Note that P15 was placed at the leftmost branch of the phylogenetic tree, which made the sequence P15 distinguishable from the rest of the T-ORF8 variants.

The pairs of T-ORF8 variants (P13, P14), (P5, P6), (P33, P45), (P9, P37), (P19, P20), (P35, P36), (P31, P34), (P29, P43), (P27, P42), (P25, P26), (P22, P23), (P21, P40), (P17, P18), (P2, P8), and (P1, P47) were found to be the closest enough to each other based on the amino acid sequence homology-based phylogeny (Figure 4).

### 3.2. Evaluation of intrinsic disorder content of 4-7 T-ORF8 proteins

We also analyzed the peculiarities of the distribution of per-residue intrinsic disorder predisposition within sequences of 47 T-ORF8 variants. Since the amino acid sequences of T-ORF8 proteins are shorter than 30 residues, the number of computational tool capable of prediction of intrinsic disorder is limited.

In this study, we used PONDR-VSL2 algorithm. Results of this analysis are shown in Figure 5. Due to their short length and limited sequence variability, T-ORF8 proteins are characterized by rather feature-less disorder profiles, where both N- and C-terminal regions are predicted to have higher levels of intrinsic disorder than the central parts.

**Figure 5:**
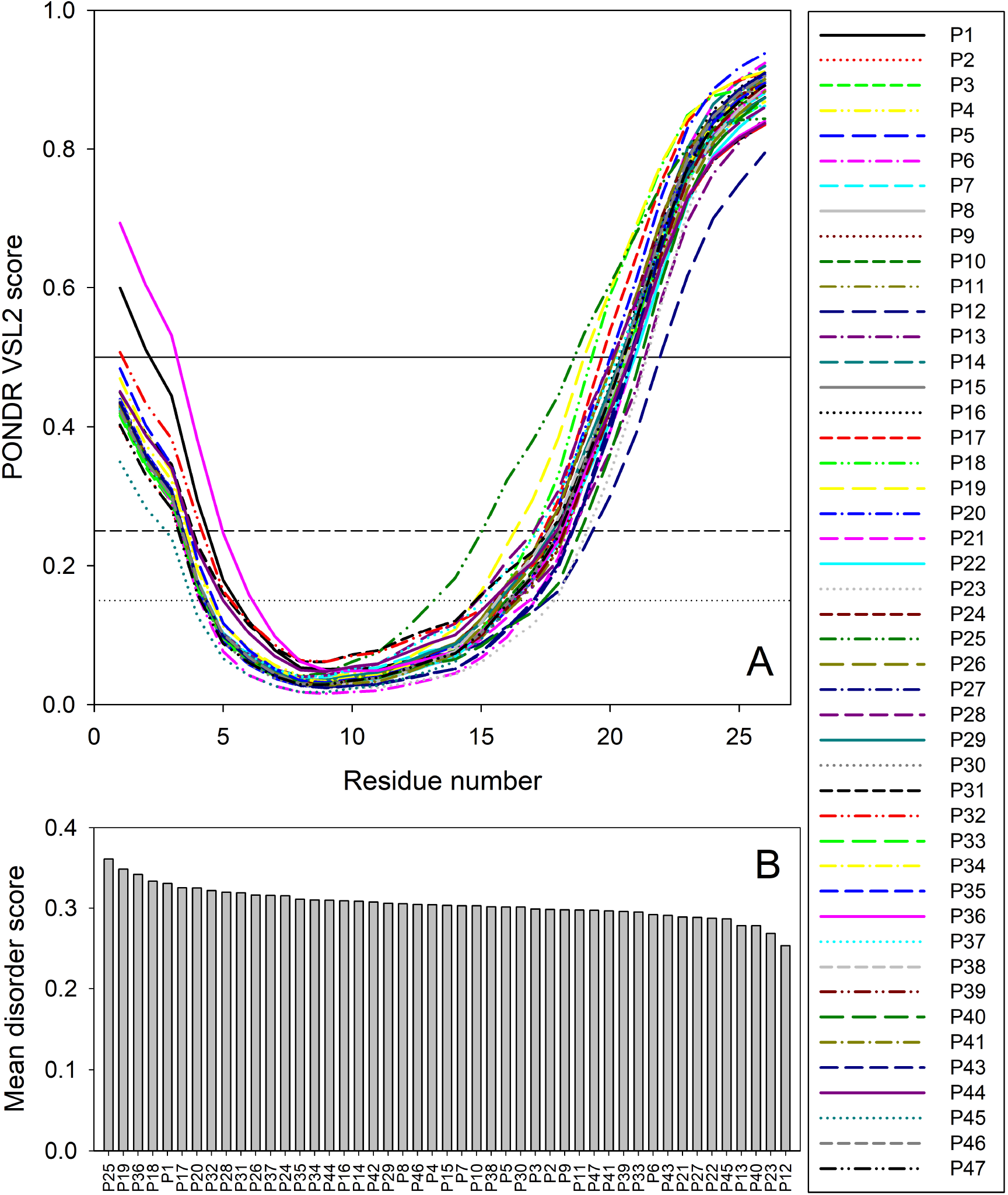
Analysis of intrinsic disorder predisposition of 47 T-ORF8 proteins: (A) Disorder profiles generated using the PONDR-VSL2 disorder predictor. Three thresholds of predicted disorder scores (PDSs) are shown, 0.15, 0.25, and 0.5, which are used for the classification of protein residues as highly disordered (*PDS* ≥ 0.5), flexible (0.25 ≤ *PDS* < 0.5), moderately flexible (0.15 ≤ *PDS* < 0.25) and mostly ordered (*PDS* < 0.15). (B) Ranking 47 T-ORF8 proteins based on their mean disorder scores.

Most T-ORF8 proteins show rather similar profiles, with the noticeable exceptions to P1 and P36 that show highest disorder levels in their N-terminal regions, P45 with least disorder N-tail, P25 with longest and most peculiar disorder distribution in its C-terminal half, P18 and P19 with long disorder stretches in their C-tails, and P12 with least levels of disorder in C-terminal regions (see Figure 5(A)). These observations are further supported by Figure 5(B), where 47 T-ORF8 proteins are ranked based on their mean disorder scores, from highest to lowest levels of disorder. Although the vast majority of T-ORF8 proteins (38 of 47) form a rather uniform cluster with the average mean disorder score of 0.304 ± 0.010, whereas P25, P19, P36, P18, and P1 showing higher than average and P13, P40, P23, and P12 lower than average levels of disorder.

**Supplementary Table S5** lists potential functional motifs identified in 47 T-ORF8 variants by ELM resource and shows that all these proteins have several such motifs. Based on the their content of functional motifs, T-ORF8 proteins can be grouped into 21 clusters, with three clusters containing 13, 4, and 2 proteins, and all the remaining being singletons. The common motif found in all T-ORF8 proteins is the N-degron that initiates protein degradation by binding to the UBR-box of N-recognins. A kinase docking motif that mediates interaction towards the ERK1/2 and P38 subfamilies of MAP kinases and a Ser/Thr residue phosphorylated by the Plk1 kinase are present in 20 clusters, whereas 17 clusters also include a site for attachment of a fucose residue to a serine. Lowest number of functional motifs (3) is found in 6 proteins (P12, P16, P17, P19, P21, and P40), many of which are characterized by lower mean disorder scores. On the contrary, proteins with largest number of functional motifs (6 and 7) are typically on a side with higher disorder scores. **Supplementary Table S5** shows that truncation might generate functional T-ORF8 variants (or at least variants possessing functional motifs), and that expected functionality of different T-ORF8 proteins can be quite different. It is clear that the results of this computational analysis should be taken with caution, and functionality of T-ORF8 requires experimental validation.

### 3.3. Variability and commonality of T-ORF8 variants

In the proceeding section, unique T-ORF8 variants were quantified using various parameters such as polar/non-polar residue sequence homology, amino acid frequency distributions, amino acid conservation through the Shannon entropy, and physicochemical properties.

#### 3.3.1. Polarity based variability of T-ORF8 variants

Each unique T-ORF8 variant possessed a binary polar/non-polar sequence and based on the sequence homology of these sequences, a phylogenetic relationship has been obtained (Figure 6).

**Figure 6:**
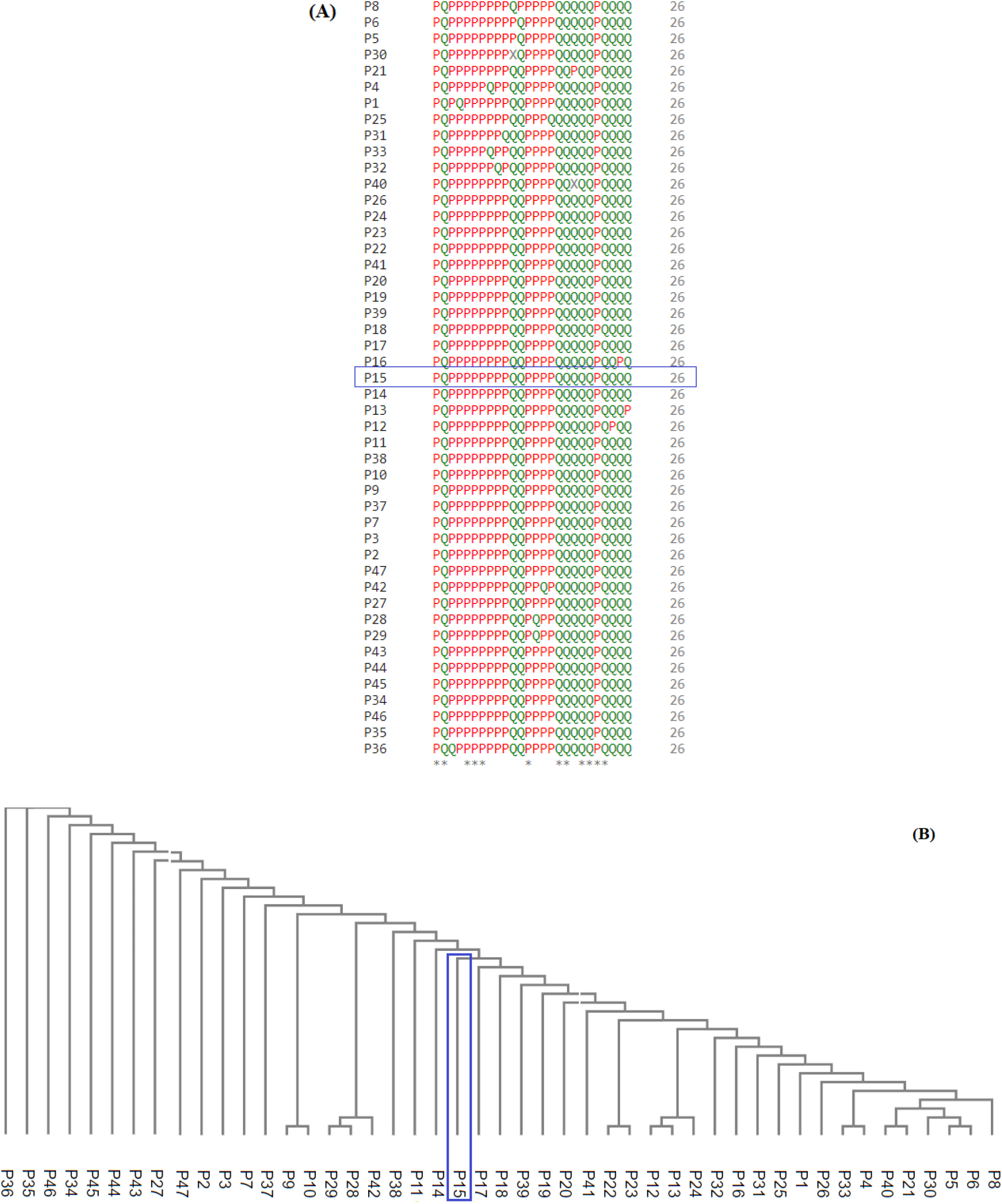
Analysis of variability among unique T-ORF8 variants based on polarity: (A) Polarity sequence-based alignment of unique T-ORF8 proteins using Clustal-Omega, and (B) associated phylogenetic relationship among the unique T-ORF8 proteins.

The number of polar and non-polar residues in the unique T-ORF8 variants was found to be almost balanced (50-50 in percentage). Among 26 residue positions of each T-ORF8 variants of amino acid length 26 residues at 14 positions (Polar residues at the positions 1, 5-7, 13 and non-polar residues at the positions 2, 17-18, 20-23) remained invariant as observed in Figure 6. The pairs of unique T-ORF8 variants (P5, P6), (P21, P40), (P4, P33), (P12, P13), (P28, P29), and (P9, P10) were closest to each other (Figure 6). Note that, the P15 variant was placed in a single leaf and found to be distant from the other unique ORF8 variants as per polarity-based homology, although P15 was found to be the closest to the T-ORF8 variants P13 and P14 based on amino acid homology.

Furthermore, it was noticed that only 17 unique T-ORF8 variants possessed unique polar/non-polar sequences (Table 7).

**Table 7:**
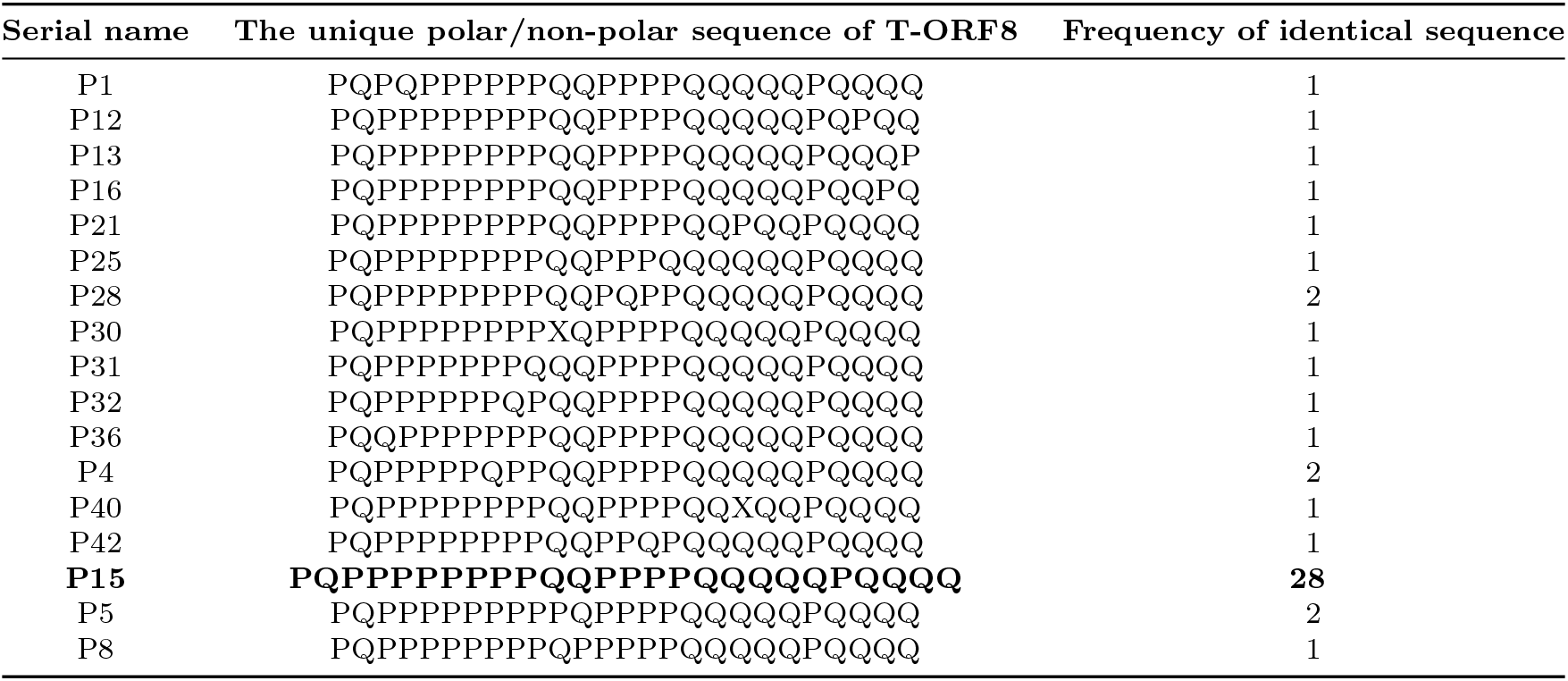
Unique polar/non-polar sequence variants of T-ORF8 and their frequencies

The polar/non-polar sequence of each T-ORF8 variant other than P4, P5, P15, and P28 was unique. Surprisingly, among the total of 47 T-ORF8 variants, there were twenty-eight T-ORF8 variants, which share identical polar/non-polar sequence with that of P15.

According to the phylogenetic relationship derived from the unique polar/non-polar sequence homology, the T-ORF8 P15 was found to be the closest to P42. Furthermore, the pairs (P13, P25), (P21, P40), (P5, P30) were found to be close enough to each other (Figure 7).

**Figure 7:**
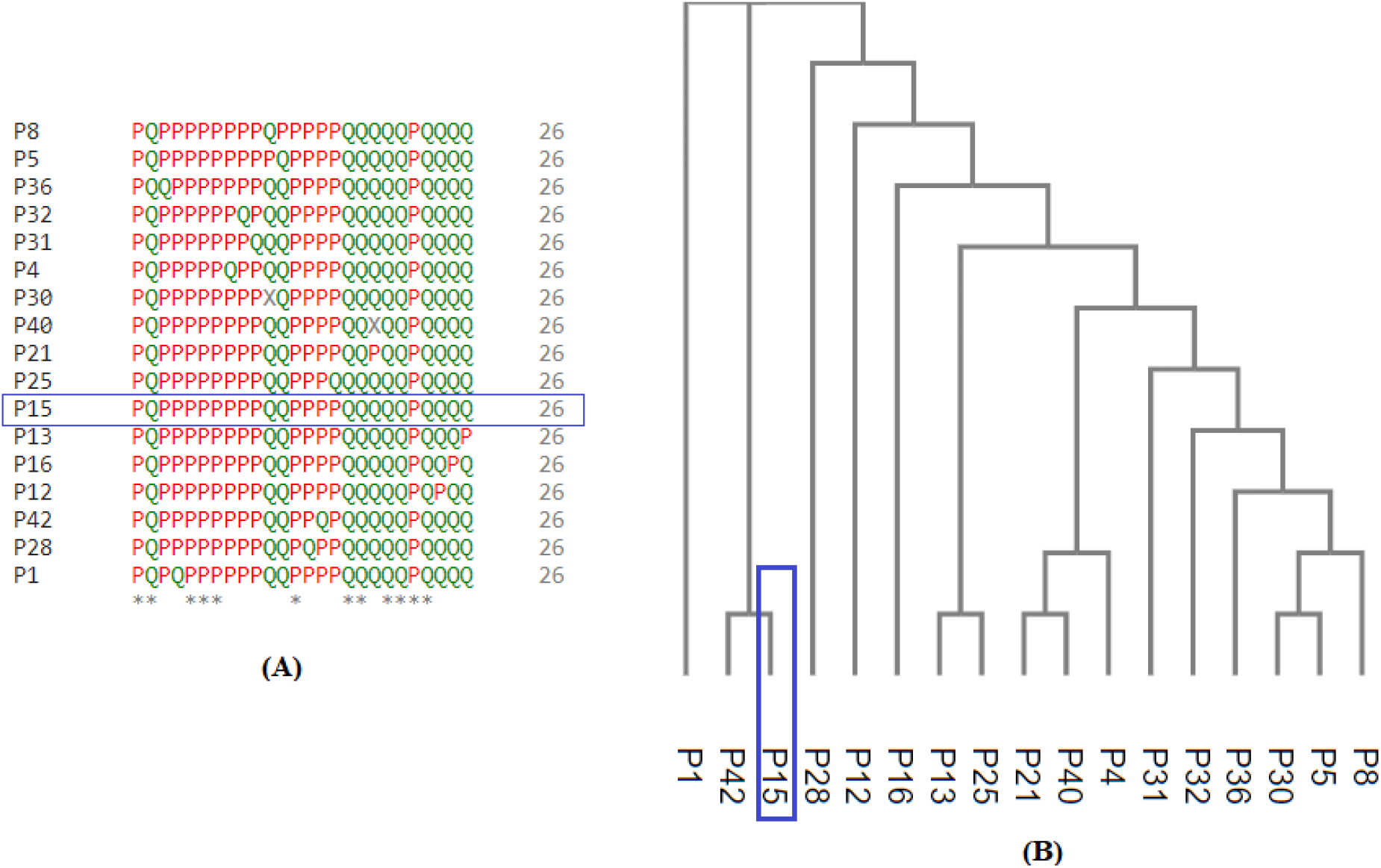
Unique polar/non-polar sequence based alignment of unique T-ORF8 proteins using the Clustal-Omega and associated phylogenetic relationship. (A) multiple sequence alignments of unique T-ORF8, and (B) phylogenetic analysis of the same samples.

#### 3.3.2. Variability of the frequency distribution of amino acids present in T-ORF8 variants

The frequency of each amino acid present in the unique T-ORF8 variants was enumerated, and consequently, a twenty-dimensional frequency vector was obtained (Table 8).

**Table 8:** Frequency of amino acids present in the 47 unique T-ORF8 variants. (Standard single-letter amino acid codes were used.)

|  | A | R | N | D | C | Q | E | G | H | I | L | K | M | F | P | S | T | W | Y | V |
| --- | --- | --- | --- | --- | --- | --- | --- | --- | --- | --- | --- | --- | --- | --- | --- | --- | --- | --- | --- | --- |
| P47 | 2 | 0 | 0 | 0 | 2 | 2 | 1 | 1 | 1 | 2 | 2 | 1 | 1 | 4 | 0 | 2 | 3 | 0 | 0 | 2 |
| P1 | 2 | 0 | 0 | 0 | 2 | 2 | 1 | 1 | 2 | 2 | 2 | 1 | 1 | 3 | 0 | 2 | 3 | 0 | 0 | 2 |
| P2 | 2 | 0 | 0 | 0 | 2 | 2 | 1 | 1 | 1 | 3 | 3 | 1 | 1 | 3 | 0 | 2 | 3 | 0 | 0 | 1 |
| P3 | 2 | 0 | 0 | 0 | 2 | 2 | 1 | 1 | 1 | 2 | 2 | 1 | 1 | 4 | 0 | 2 | 3 | 0 | 0 | 2 |
| P4 | 2 | 0 | 0 | 0 | 2 | 2 | 2 | 0 | 1 | 2 | 3 | 1 | 1 | 3 | 0 | 2 | 3 | 0 | 0 | 2 |
| P5 | 3 | 0 | 0 | 0 | 2 | 2 | 1 | 1 | 1 | 2 | 3 | 1 | 1 | 3 | 0 | 2 | 2 | 0 | 0 | 2 |
| P6 | 2 | 0 | 0 | 0 | 2 | 2 | 1 | 1 | 1 | 3 | 3 | 1 | 1 | 3 | 0 | 2 | 2 | 0 | 0 | 2 |
| P7 | 2 | 0 | 0 | 0 | 2 | 2 | 1 | 1 | 1 | 2 | 3 | 2 | 1 | 3 | 0 | 2 | 2 | 0 | 0 | 2 |
| P8 | 2 | 0 | 0 | 0 | 2 | 2 | 1 | 1 | 1 | 2 | 3 | 1 | 1 | 3 | 1 | 2 | 2 | 0 | 0 | 2 |
| P37 | 3 | 0 | 0 | 0 | 2 | 2 | 1 | 1 | 1 | 2 | 3 | 1 | 1 | 3 | 0 | 2 | 3 | 0 | 0 | 1 |
| P9 | 2 | 0 | 0 | 0 | 2 | 2 | 1 | 1 | 1 | 3 | 3 | 1 | 1 | 3 | 0 | 2 | 3 | 0 | 0 | 1 |
| P10 | 2 | 0 | 0 | 0 | 2 | 2 | 1 | 1 | 1 | 2 | 4 | 1 | 1 | 3 | 0 | 2 | 3 | 0 | 0 | 1 |
| P38 | 2 | 0 | 0 | 1 | 2 | 2 | 1 | 1 | 0 | 2 | 3 | 1 | 1 | 3 | 0 | 2 | 3 | 0 | 0 | 2 |
| P11 | 2 | 0 | 0 | 1 | 2 | 2 | 0 | 1 | 1 | 2 | 3 | 1 | 1 | 3 | 0 | 2 | 3 | 0 | 0 | 2 |
| P12 | 2 | 0 | 0 | 0 | 2 | 2 | 1 | 1 | 1 | 2 | 4 | 1 | 1 | 3 | 0 | 1 | 3 | 0 | 0 | 2 |
| P13 | 2 | 0 | 0 | 0 | 2 | 2 | 1 | 1 | 1 | 3 | 3 | 1 | 1 | 3 | 0 | 2 | 2 | 0 | 0 | 2 |
| P14 | 2 | 0 | 1 | 0 | 2 | 2 | 1 | 1 | 1 | 2 | 3 | 1 | 1 | 3 | 0 | 2 | 2 | 0 | 0 | 2 |
| P15 | 2 | 0 | 0 | 0 | 2 | 2 | 1 | 1 | 1 | 2 | 3 | 1 | 1 | 3 | 0 | 2 | 3 | 0 | 0 | 2 |
| P16 | 2 | 0 | 0 | 0 | 1 | 2 | 1 | 1 | 1 | 2 | 3 | 1 | 1 | 4 | 0 | 2 | 3 | 0 | 0 | 2 |
| P17 | 2 | 1 | 0 | 0 | 1 | 2 | 1 | 1 | 1 | 2 | 3 | 1 | 1 | 3 | 0 | 2 | 3 | 0 | 0 | 2 |
| P18 | 2 | 0 | 0 | 0 | 1 | 2 | 1 | 1 | 1 | 2 | 3 | 1 | 1 | 3 | 0 | 3 | 3 | 0 | 0 | 2 |
| P39 | 2 | 1 | 0 | 0 | 2 | 1 | 1 | 1 | 1 | 2 | 3 | 1 | 1 | 3 | 0 | 2 | 3 | 0 | 0 | 2 |
| P19 | 2 | 1 | 0 | 0 | 1 | 2 | 1 | 1 | 1 | 2 | 3 | 1 | 1 | 3 | 0 | 2 | 3 | 0 | 0 | 2 |
| P20 | 2 | 0 | 0 | 0 | 1 | 2 | 1 | 1 | 1 | 2 | 3 | 1 | 1 | 3 | 0 | 2 | 3 | 0 | 1 | 2 |
| P21 | 2 | 0 | 0 | 0 | 2 | 2 | 0 | 2 | 1 | 2 | 3 | 1 | 1 | 3 | 0 | 2 | 3 | 0 | 0 | 2 |
| P40 | 2 | 0 | 0 | 0 | 2 | 2 | 0 | 1 | 1 | 2 | 3 | 1 | 1 | 3 | 0 | 2 | 3 | 0 | 0 | 2 |
| P41 | 2 | 0 | 1 | 0 | 2 | 2 | 1 | 1 | 0 | 2 | 3 | 1 | 1 | 3 | 0 | 2 | 3 | 0 | 0 | 2 |
| P22 | 2 | 1 | 0 | 0 | 2 | 2 | 1 | 1 | 0 | 2 | 3 | 1 | 1 | 3 | 0 | 2 | 3 | 0 | 0 | 2 |
| P23 | 2 | 0 | 0 | 0 | 2 | 2 | 1 | 1 | 0 | 2 | 3 | 1 | 1 | 3 | 0 | 2 | 3 | 0 | 1 | 2 |
| P24 | 2 | 0 | 0 | 0 | 2 | 2 | 1 | 1 | 1 | 2 | 4 | 1 | 1 | 2 | 0 | 2 | 3 | 0 | 0 | 2 |
| P25 | 2 | 0 | 0 | 0 | 2 | 2 | 1 | 1 | 1 | 2 | 3 | 1 | 1 | 2 | 0 | 3 | 3 | 0 | 0 | 2 |
| P26 | 2 | 0 | 0 | 0 | 2 | 2 | 1 | 1 | 1 | 2 | 3 | 1 | 1 | 2 | 0 | 2 | 3 | 0 | 0 | 3 |
| P42 | 1 | 0 | 0 | 0 | 2 | 2 | 1 | 1 | 1 | 2 | 3 | 1 | 1 | 3 | 0 | 2 | 4 | 0 | 0 | 2 |
| P27 | 1 | 0 | 0 | 0 | 2 | 2 | 1 | 1 | 1 | 2 | 3 | 1 | 1 | 3 | 0 | 2 | 3 | 0 | 0 | 3 |
| P28 | 1 | 0 | 0 | 0 | 2 | 2 | 1 | 1 | 1 | 2 | 3 | 1 | 1 | 3 | 0 | 3 | 3 | 0 | 0 | 2 |
| P29 | 1 | 0 | 0 | 0 | 2 | 2 | 1 | 1 | 1 | 2 | 3 | 1 | 1 | 3 | 0 | 2 | 4 | 0 | 0 | 2 |
| P43 | 1 | 0 | 0 | 0 | 2 | 2 | 1 | 1 | 1 | 2 | 3 | 1 | 1 | 3 | 0 | 2 | 3 | 0 | 0 | 3 |
| P30 | 2 | 0 | 0 | 0 | 2 | 2 | 1 | 1 | 1 | 2 | 3 | 1 | 1 | 3 | 0 | 2 | 2 | 0 | 0 | 2 |
| P44 | 2 | 0 | 0 | 0 | 2 | 2 | 1 | 1 | 1 | 1 | 3 | 1 | 2 | 3 | 0 | 2 | 3 | 0 | 0 | 2 |
| P31 | 2 | 0 | 0 | 0 | 2 | 2 | 1 | 1 | 1 | 1 | 3 | 1 | 1 | 3 | 0 | 2 | 4 | 0 | 0 | 2 |
| P32 | 2 | 0 | 0 | 0 | 2 | 2 | 1 | 1 | 1 | 1 | 3 | 1 | 1 | 3 | 0 | 2 | 4 | 0 | 0 | 2 |
| P33 | 2 | 1 | 0 | 0 | 2 | 2 | 1 | 0 | 1 | 2 | 3 | 1 | 1 | 3 | 0 | 2 | 3 | 0 | 0 | 2 |
| P45 | 2 | 0 | 0 | 0 | 2 | 2 | 1 | 0 | 1 | 2 | 3 | 1 | 1 | 3 | 0 | 2 | 3 | 0 | 0 | 3 |
| P34 | 2 | 0 | 0 | 0 | 2 | 2 | 1 | 1 | 1 | 2 | 4 | 1 | 1 | 2 | 0 | 2 | 3 | 0 | 0 | 2 |
| P46 | 2 | 0 | 0 | 0 | 2 | 2 | 1 | 1 | 1 | 3 | 3 | 1 | 1 | 2 | 0 | 2 | 3 | 0 | 0 | 2 |
| P35 | 2 | 0 | 0 | 0 | 2 | 2 | 1 | 1 | 1 | 2 | 4 | 1 | 1 | 2 | 0 | 2 | 3 | 0 | 0 | 2 |
| P36 | 2 | 0 | 0 | 0 | 2 | 2 | 1 | 1 | 1 | 2 | 3 | 1 | 1 | 2 | 0 | 3 | 3 | 0 | 0 | 2 |

Tryptophan was not present in any of the unique T-ORF8 variants. It was noted that the amino acids arginine, asparagine, aspartic acid, proline and tyrosine were absent in the T-ORF8 P15. The amino acid arginine was found with frequency one in the T-ORF8 P17, P39, P19, P22 and P33. In the sequence P14 and P41, asparagine was present with frequency one. Likewise, aspartic acid was found in P38 and P11. The amino acid proline with frequency one was found in the P8 variant only. In the T-ORF8 variants, P20 and P23 tyrosine was found. The highest frequency of each amino acids phenylalanine (in P3, P16, and P47), leucine (in P10, P12, P24, P34, and P35), and threonine (in P29, P31, P32, and P42) were 4.

For each pair of frequency vectors corresponding to all the unique T-ORF8 variants, Euclidean distances were calculated (*Supplementary file-1*), and the distance matrix in color heat-map is presented in Figure 8. It was found that the P15 variant is equidistant (1.41) from all other variants except P30 and P40 which were 1 distance apart from P15.

**Figure 8:**
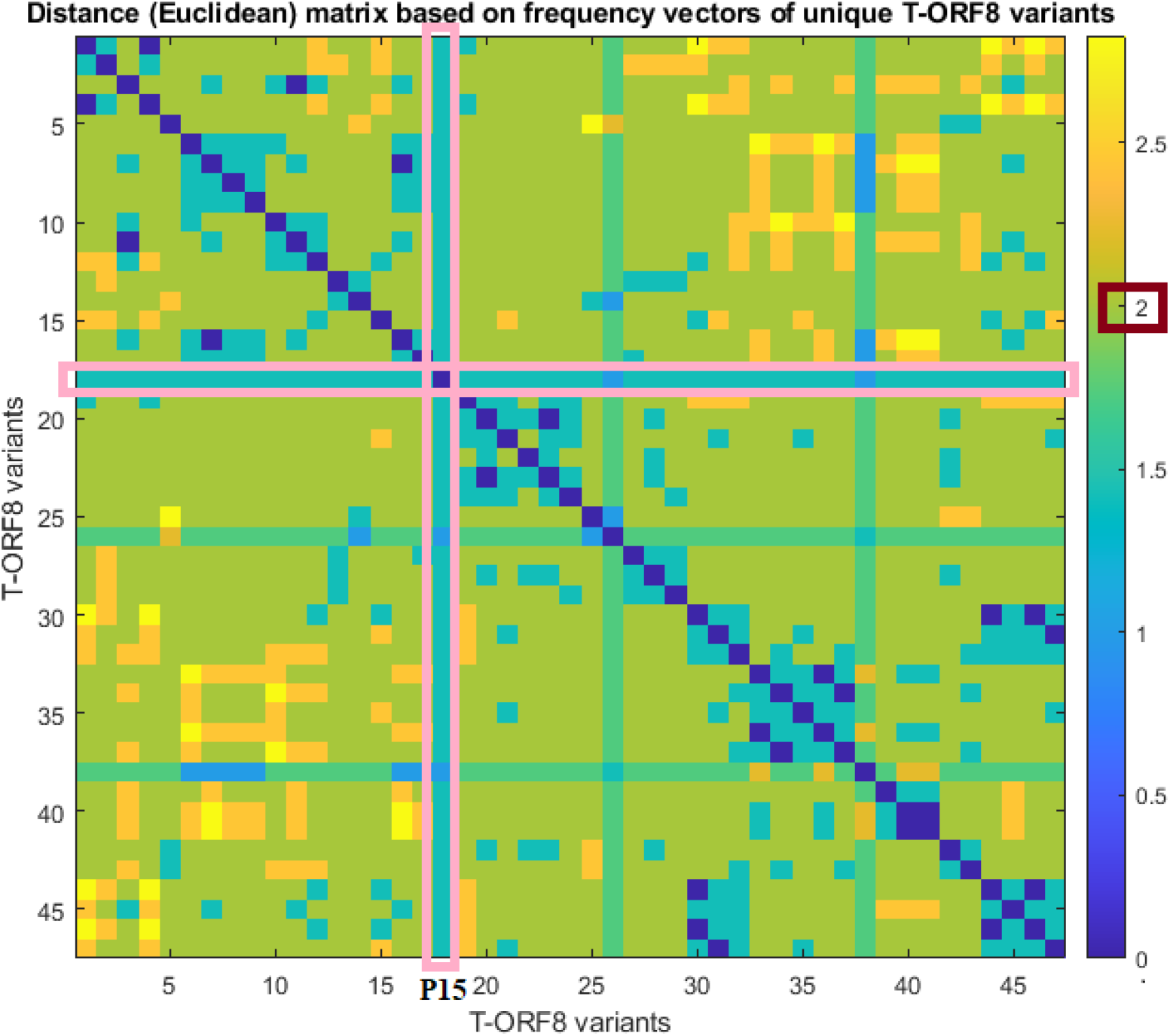
Pairwise distance matrix of amino acid frequency vectors of the unique T-ORF8 variants.

Further, we observed that the distance between any two pairs of T-ORF8 variants is 2 (light green color) except for a few cases (Figure 7). Although the amino acid sequences were different, identical frequency vectors were found for the pair of ORF8 variants (P3, P47), (P2, P9), (P6, P13), (P17, P19), (P24, P34), (P24, P35), (P25, P36), (P34, P35), (P29, P42), and (P27, P43).

Based on the distance matrix, all the unique T-ORF8 variants were clustered, and the associated phylogeny is presented in Figure 8. The P15 variant was very close to P22, P23, P33, P40, and P41 according to the phylogenetic relationship depicted in Figure 9.

**Figure 9:**
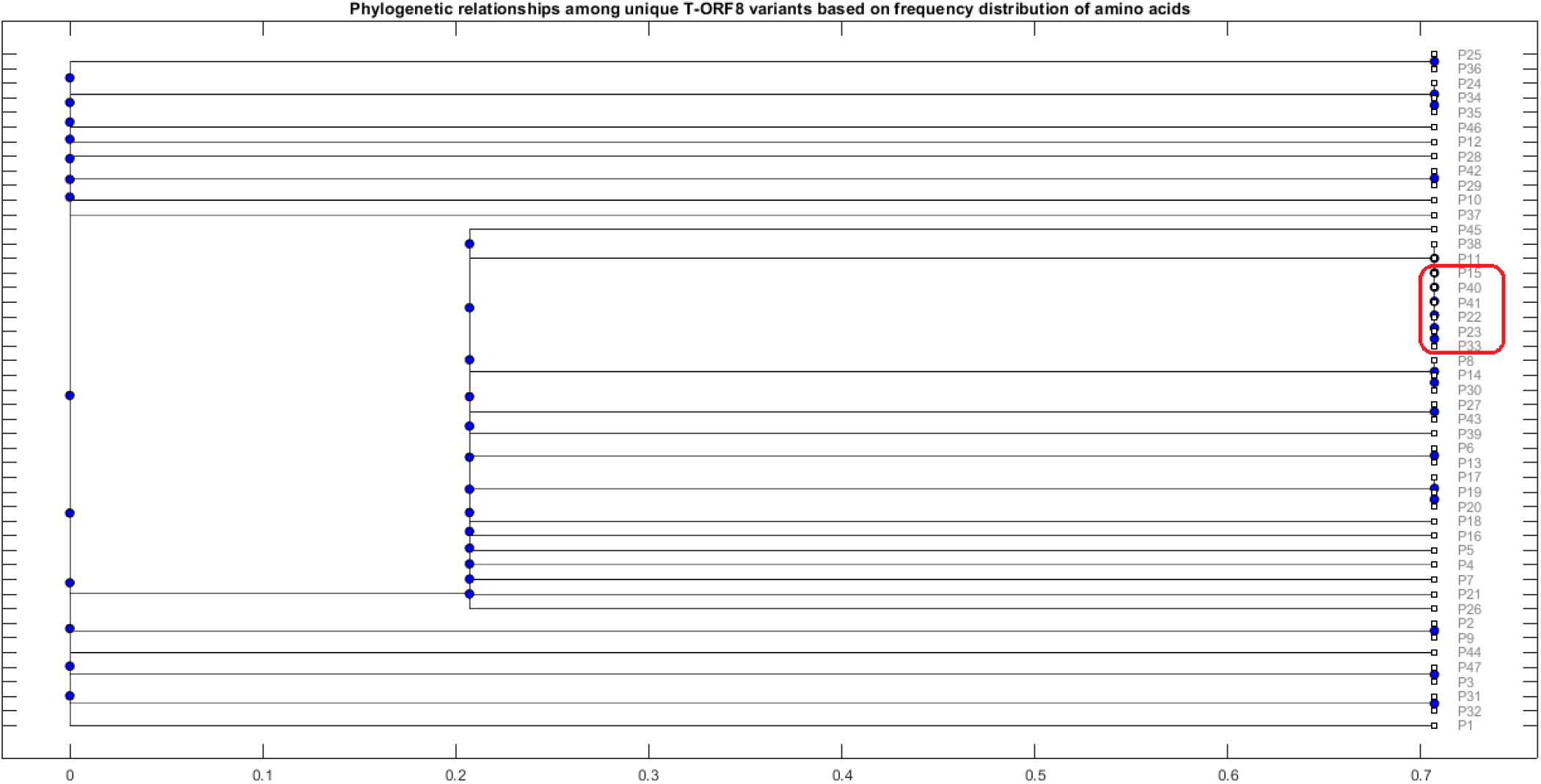
Frequency distribution of amino acids present in the unique T-ORF8 variants and their phylogenetic relationships. The red box highlights the closeness of P15 with P22, P23, P33, P40, and P41 variants.

Other than the pairs of T-ORF8 having identical frequency vectors, it was found that the pairs of unique T-ORF8 variants (P23, P33), (P14, P30), (P19, P20), and (P31, P32) were close to each other as derived from the phylogenetic relationship (Figure 9).

#### 3.3.3. Variability of T-ORF8 through Shannon entropy

Shannon entropy for each unique T-ORF8 variant was calculated using the formula stated in section 2.3 (Table 9). It was found that the highest and lowest SEs of 47 unique T-ORF8 proteins were 0.958 and 0.973 respectively. That is, the length of the largest interval is 0.015, which is sufficiently small. Based on SEs of the T-ORF8 proteins a set of clusters were derived (Figure 10 (A)), and SEs of each of the T-ORF8 variants are plotted in Figure 10 (B).

**Figure 10:**
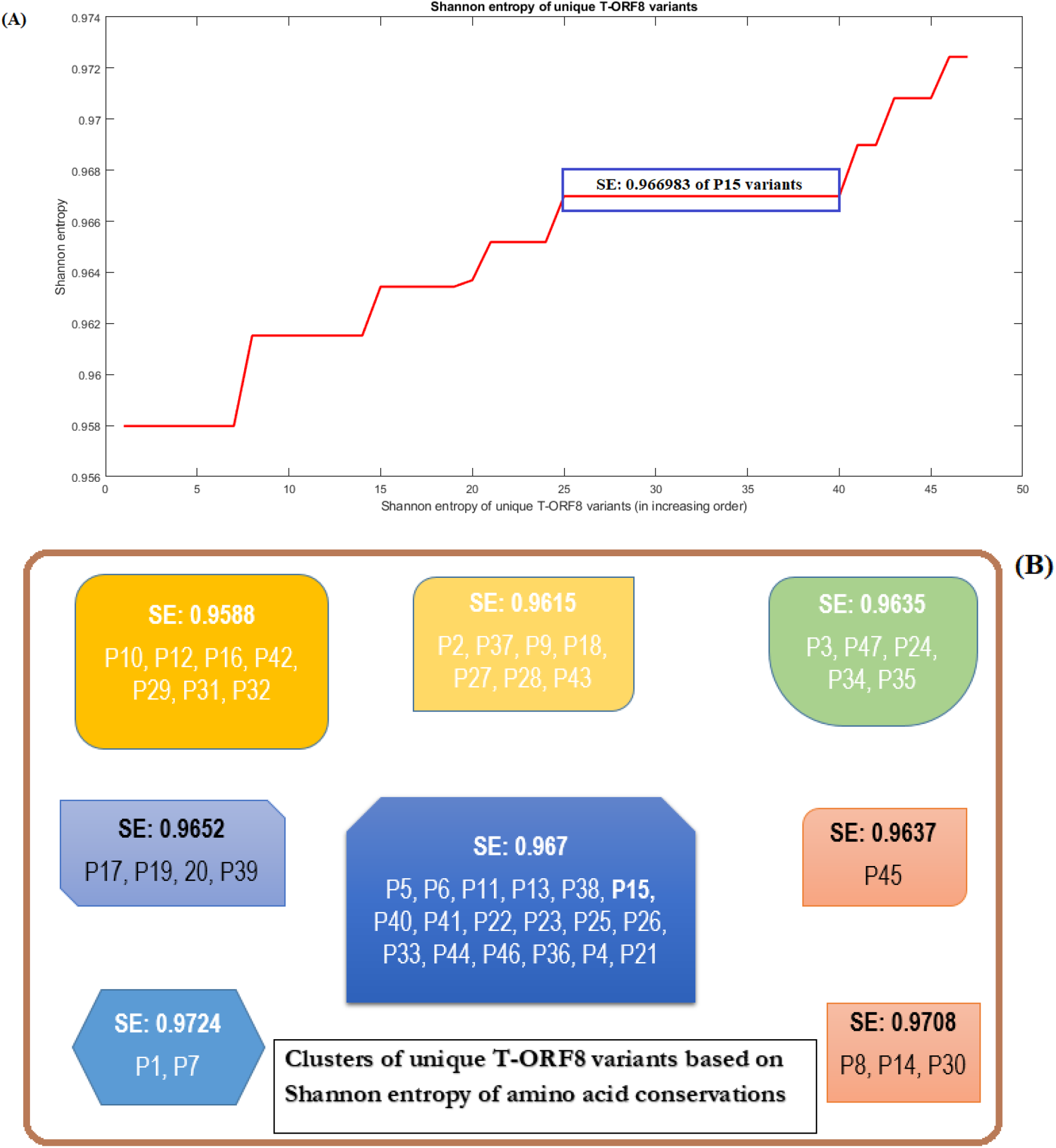
Analysis of T-ORF8 variability:(A) Shannon entropy of unique T-ORF8 protein variants (plotted in increasing order), and (B) clusters of T-ORF8 variants based on Shannon entropy. The boxes highlighted in different colors represent clusters of unique T-ORF8 variants.

**Table 9:** Shannon entropy of amino acid conservations over the unique T-ORF8 variants

| Serial name | SE | Serial name | SE | Serial name | SE |
| --- | --- | --- | --- | --- | --- |
| P10 | 0.957979735 | P24 | 0.963438 | P22 | 0.966983 |
| P12 | 0.957979735 | P34 | 0.963438 | P23 | 0.966983 |
| P16 | 0.957979735 | P35 | 0.963438 | P25 | 0.966983 |
| P42 | 0.957979735 | P45 | 0.963693 | P26 | 0.966983 |
| P29 | 0.957979735 | P17 | 0.965186 | P44 | 0.966983 |
| P31 | 0.957979735 | P39 | 0.965186 | P33 | 0.966983 |
| P32 | 0.957979735 | P19 | 0.965186 | P46 | 0.966983 |
| P2 | 0.961524341 | P20 | 0.965186 | P36 | 0.966983 |
| P37 | 0.961524341 | P5 | 0.966983 | P4 | 0.968986 |
| P9 | 0.961524341 | P6 | 0.966983 | P21 | 0.968986 |
| P18 | 0.961524341 | P38 | 0.966983 | P8 | 0.970821 |
| P27 | 0.961524341 | P11 | 0.966983 | P14 | 0.970821 |
| P28 | 0.961524341 | P13 | 0.966983 | P30 | 0.970821 |
| P43 | 0.961524341 | **P15** | **0.966983** | P1 | 0.972441 |
| P47 | 0.963438001 | P40 | 0.966983 | P7 | 0.972441 |
| P3 | 0.963438001 | P41 | 0.966983 |  |  |

The largest cluster containing 18 (among which the T-ORF8 P15 variant also present) T-ORF8 variants based on the identical SEs were obtained (Figure 10).

#### 3.3.4. Molecular and physicochemical informatics of T-ORF8 unique variants

For each unique T-ORF8 variant and complete ORF8 protein, several physicochemical and molecular properties were computed using the web-servers as mentioned in section 2.4 (Table 10). It was found that the extinction coefficient of all the T-ORF8 variants was found to be 125, except for four T-ORF8 variants P16, P17, P18, and P19 whose extinction coefficient was zero (Table 9). Further, it was noticed that for P20 and P23 extinction coefficients were found to be significantly high compared to others. Instability indices of all the T-ORF8 protein variants were ranging from 45.36 to 95.85 (greater than 40), and consequently they all are unstable. It was observed that the P15 variants had a unique frequency of the various type of residues (Tiny: 10, Small: 12, Aliphatic: 9, Aromatic: 4, Non-polar: 16, Polar: 10, Charged: 3, Basic: 2, Acidic: 1) and none of the other T-ORF8 variants had it identical.

**Table 10:** Molecular and physicochemical informatics of T-ORF8 unique variants

| Sl. Name | PI | EC | II | AI | GRAVY | PS | Tiny | Small | Aliphatic | Aromatic | Non-polar | Polar | Charged | Basic | Acidic |
| --- | --- | --- | --- | --- | --- | --- | --- | --- | --- | --- | --- | --- | --- | --- | --- |
| ORF8 | 5.42 | 16305 | 45.36 | 97.36 | 0.219 | 0.462 | 31 | 59 | 37 | 20 | 73 | 48 | 26 | 13 | 13 |
| P1 | 6.68 | 125 | 85.17 | 90 | 0.731 | 0.553 | 10 | 12 | 8 | 5 | 15 | 11 | 4 | 3 | 1 |
| P2 | 6.49 | 125 | 85.17 | 108.85 | 1.012 | 0.436 | 10 | 11 | 9 | 4 | 16 | 10 | 3 | 2 | 1 |
| P3 | 6.49 | 125 | 85.17 | 90 | 0.962 | 0.553 | 10 | 12 | 8 | 5 | 16 | 10 | 3 | 2 | 1 |
| P4 | 5.38 | 125 | 95.85 | 105 | 0.881 | 0.578 | 9 | 11 | 9 | 4 | 15 | 11 | 4 | 2 | 2 |
| P5 | 6.49 | 125 | 85.17 | 108.85 | 1.096 | 0.492 | 10 | 12 | 10 | 4 | 17 | 9 | 3 | 2 | 1 |
| P6 | 6.49 | 125 | 85.17 | 120 | 1.2 | 0.492 | 9 | 11 | 10 | 4 | 17 | 9 | 3 | 2 | 1 |
| P7 | 7.84 | 125 | 81.91 | 105 | 0.877 | 0.578 | 9 | 11 | 9 | 4 | 16 | 10 | 4 | 3 | 1 |
| P8 | 6.49 | 125 | 92.58 | 105 | 0.965 | 0.492 | 9 | 12 | 9 | 4 | 17 | 9 | 3 | 2 | 1 |
| P9 | 6.49 | 125 | 85.17 | 108.85 | 1.012 | 0.436 | 10 | 11 | 9 | 4 | 16 | 10 | 3 | 2 | 1 |
| P10 | 6.49 | 125 | 85.17 | 108.85 | 0.985 | 0.436 | 10 | 11 | 9 | 4 | 16 | 10 | 3 | 2 | 1 |
| P11 | 6.49 | 125 | 68.27 | 105 | 1 | 0.492 | 10 | 13 | 9 | 4 | 16 | 10 | 3 | 2 | 1 |
| P12 | 6.49 | 125 | 55.73 | 120 | 1.177 | 0.492 | 9 | 11 | 10 | 4 | 17 | 9 | 3 | 2 | 1 |
| P13 | 6.49 | 125 | 72.63 | 120 | 1.2 | 0.492 | 9 | 11 | 10 | 4 | 17 | 9 | 3 | 2 | 1 |
| P14 | 6.49 | 125 | 72.63 | 105 | 0.892 | 0.492 | 9 | 12 | 9 | 4 | 16 | 10 | 3 | 2 | 1 |
| P15 | 6.49 | 125 | 85.17 | 105 | 1 | 0.492 | 10 | 12 | 9 | 4 | 16 | 10 | 3 | 2 | 1 |
| P16 | 6.5 | 0 | 60.1 | 105 | 1.012 | 0.492 | 9 | 11 | 9 | 5 | 16 | 10 | 3 | 2 | 1 |
| P17 | 8 | 0 | 67.5 | 105 | 0.731 | 0.544 | 9 | 11 | 9 | 4 | 15 | 11 | 4 | 3 | 1 |
| P18 | 6.5 | 0 | 67.5 | 105 | 0.873 | 0.492 | 10 | 12 | 9 | 4 | 15 | 11 | 3 | 2 | 1 |
| P19 | 8 | 0 | 85.17 | 105 | 0.731 | 0.544 | 9 | 11 | 9 | 4 | 15 | 11 | 4 | 3 | 1 |
| P21 | 7.85 | 125 | 60.87 | 105 | 1.119 | 0.578 | 11 | 13 | 9 | 4 | 17 | 9 | 2 | 2 | 0 |
| P22 | 7.82 | 125 | 92.58 | 105 | 0.95 | 0.597 | 10 | 12 | 9 | 3 | 16 | 10 | 3 | 2 | 1 |
| P24 | 6.49 | 125 | 85.17 | 120 | 1.038 | 0.543 | 10 | 12 | 10 | 3 | 16 | 10 | 3 | 2 | 1 |
| P25 | 6.49 | 125 | 85.17 | 105 | 0.862 | 0.543 | 11 | 13 | 9 | 3 | 15 | 11 | 3 | 2 | 1 |
| P26 | 6.49 | 125 | 85.17 | 116.15 | 1.054 | 0.543 | 10 | 13 | 10 | 3 | 16 | 10 | 3 | 2 | 1 |
| P27 | 6.49 | 125 | 85.17 | 112.31 | 1.092 | 0.492 | 9 | 12 | 9 | 4 | 16 | 10 | 3 | 2 | 1 |
| P28 | 6.49 | 125 | 85.17 | 101.15 | 0.9 | 0.492 | 10 | 12 | 8 | 4 | 15 | 11 | 3 | 2 | 1 |
| P29 | 6.49 | 125 | 81.91 | 101.15 | 0.904 | 0.492 | 10 | 12 | 8 | 4 | 15 | 11 | 3 | 2 | 1 |
| P30 | 6.49 | 125 | 84.4 | 105 | 1.027 | 0.492 | 9 | 11 | 9 | 4 | 16 | 9 | 3 | 2 | 1 |
| P31 | 6.49 | 125 | 85.17 | 90 | 0.8 | 0.492 | 11 | 13 | 8 | 4 | 15 | 11 | 3 | 2 | 1 |
| P32 | 6.49 | 125 | 85.17 | 90 | 0.8 | 0.492 | 11 | 13 | 8 | 4 | 15 | 11 | 3 | 2 | 1 |
| P33 | 7.85 | 125 | 95.85 | 105 | 0.842 | 0.544 | 9 | 11 | 9 | 4 | 15 | 11 | 4 | 3 | 1 |
| P34 | 6.49 | 125 | 85.17 | 120 | 1.038 | 0.543 | 10 | 12 | 10 | 3 | 16 | 10 | 3 | 2 | 1 |
| P35 | 6.49 | 125 | 81.91 | 120 | 1.038 | 0.543 | 10 | 12 | 10 | 3 | 16 | 10 | 3 | 2 | 1 |
| P36 | 6.49 | 125 | 85.17 | 105 | 0.862 | 0.543 | 11 | 13 | 9 | 3 | 15 | 11 | 3 | 2 | 1 |
| P37 | 6.49 | 125 | 85.17 | 97.69 | 0.908 | 0.436 | 11 | 12 | 9 | 4 | 16 | 10 | 3 | 2 | 1 |
| P38 | 4.37 | 125 | 89.92 | 105 | 0.988 | 0.63 | 10 | 13 | 9 | 3 | 16 | 10 | 3 | 2 | 1 |
| P39 | 7.85 | 125 | 80.04 | 105 | 0.962 | 0.544 | 10 | 12 | 9 | 4 | 16 | 10 | 4 | 3 | 1 |
| P40 | 7.85 | 125 | 60.1 | 105 | 1.135 | 0.578 | 10 | 12 | 9 | 4 | 16 | 9 | 2 | 2 | 0 |
| P41 | 5.75 | 125 | 82.27 | 105 | 0.988 | 0.545 | 10 | 13 | 9 | 3 | 16 | 10 | 2 | 1 | 1 |
| P42 | 6.49 | 125 | 89.92 | 101.5 | 0.904 | 0.492 | 10 | 12 | 8 | 4 | 15 | 11 | 3 | 2 | 1 |
| P43 | 6.49 | 125 | 85.17 | 112.31 | 1.092 | 0.492 | 9 | 12 | 9 | 4 | 16 | 10 | 3 | 2 | 1 |
| P44 | 6.49 | 125 | 84.07 | 90 | 0.9 | 0.492 | 10 | 12 | 8 | 4 | 16 | 10 | 3 | 2 | 1 |
| P45 | 6.49 | 125 | 88.44 | 116.15 | 1.177 | 0.492 | 9 | 12 | 10 | 4 | 16 | 10 | 3 | 2 | 1 |
| P46 | 6.49 | 125 | 89.32 | 120 | 1.065 | 0.543 | 10 | 12 | 10 | 3 | 16 | 10 | 3 | 2 | 1 |
| P47 | 6.49 | 125 | 85.17 | 90 | 0.962 | 0.553 | 10 | 12 | 8 | 5 | 16 | 10 | 3 | 2 | 1 |

Furthermore, Euclidean distances between every pair of molecular and physicochemical property vectors corresponding to each T-ORF8 variant were computed and based on the distance matrix (*Supplementary file-2*) a phylogenetic relationship was derived (Figure 11).

**Figure 11:**
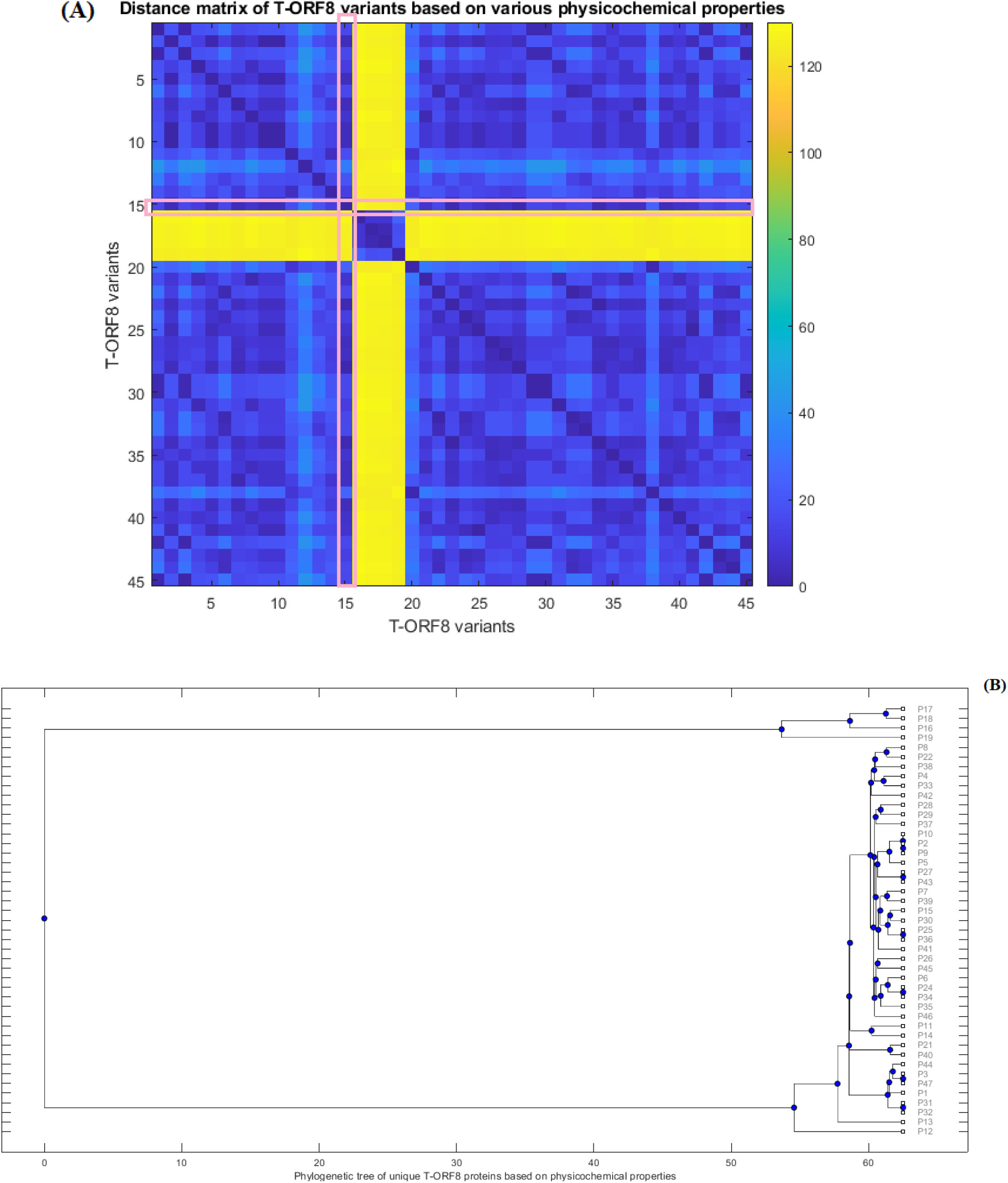
Distance matrix of property vectors and derived phylogenetic tree of 45 T-ORF8 variants. (A) represents distance matrix, (B) Phylogenetic tree based on physicochemical properties.

Note that the property vectors of P20 and P30 were highly distant from that of other ORF8 variants due to the huge difference in the extinction coefficients (for P20, EC: 1490 and for P30, EC: 1615). So ignoring these two ORF8 variants, the phylogenetic relationship among the remaining 45 T-ORF8 was derived. It was found that none of the T-ORF8 variants had identical property vectors as that of the P15 variant.

It was further found from the phylogenetic relationship that the pair of unique T-ORF8 variants (P17, P18), (P8, P22), (P4, P33), (P28, P29), (P2, P9), (P27, P43), (P7, P39), (P15, P30), (P25, P36), (P26, P45), (P24, P34), (P11, P14), (P21, P40), (P3, P47,), and (P31, P32) were found to be the closest pairs based on the closeness of property vectors.

Property vector distances from each 45 unique T-ORF8 variants from P15 are presented in Table 11.

**Table 11:** Distance from the P15 variant to the unique T-ORF8 variants, based on the physicochemical feature vectors

| Sl. Name | P15 | Sl. Name | P15 | Sl. Name | P15 | Sl. Name | P15 | Sl. Name | P15 |
| --- | --- | --- | --- | --- | --- | --- | --- | --- | --- |
| **P15** | 0 | **P5** | 4.222762 | **P22** | 7.595428 | **P3** | 15.06669 | **P11** | 16.92956 |
| **P30** | 1.895687 | **P28** | 4.222854 | **P8** | 7.609818 | **P47** | 15.06669 | **P13** | 19.67973 |
| **P25** | 2.240903 | **P29** | 5.334727 | **P4** | 11.01436 | **P44** | 15.07382 | **P21** | 24.46142 |
| **P36** | 2.240903 | **P38** | 5.392225 | **P33** | 11.04263 | **P6** | 15.16707 | **P40** | 25.16704 |
| **P41** | 3.600077 | **P39** | 5.492781 | **P26** | 11.28397 | **P31** | 15.16707 | **P12** | 33.11714 |
| **P2** | 3.978163 | **P42** | 6.149936 | **P45** | 11.7067 | **P32** | 15.16707 | **P19** | 125.0334 |
| **P9** | 3.978163 | **P27** | 7.378656 | **P14** | 12.58027 | **P1** | 15.20237 | **P18** | 126.2507 |
| **P10** | 3.978173 | **P43** | 7.378656 | **P24** | 15.06665 | **P35** | 15.41531 | **P17** | 126.2758 |
| **P7** | 4.058648 | **P37** | 7.378868 | **P34** | 15.06665 | **P46** | 15.62784 | **P16** | 127.501 |

In the close vicinity of P15, only P25, P30, and P36 variants appeared based on the nearness of property vectors (Table 10).

### 3.4. Possible T-ORF8 variants in the likelihood of P15 variant

Based on the amino acid sequence homology and other various features such as the frequency distribution of amino acids, SE, and physicochemical properties of T-ORF8 variants a possible cluster of nine unique T-ORF8 variants are derived. A schematic presentation is given in Figure 12. Note that the possible T-ORF8 variants were made of the set-theoretic union of the sets of possible T-ORF8 variants which were placed in the likelihood of P15 based on various quantitative measures mentioned in the result subsections.

**Figure 12:**
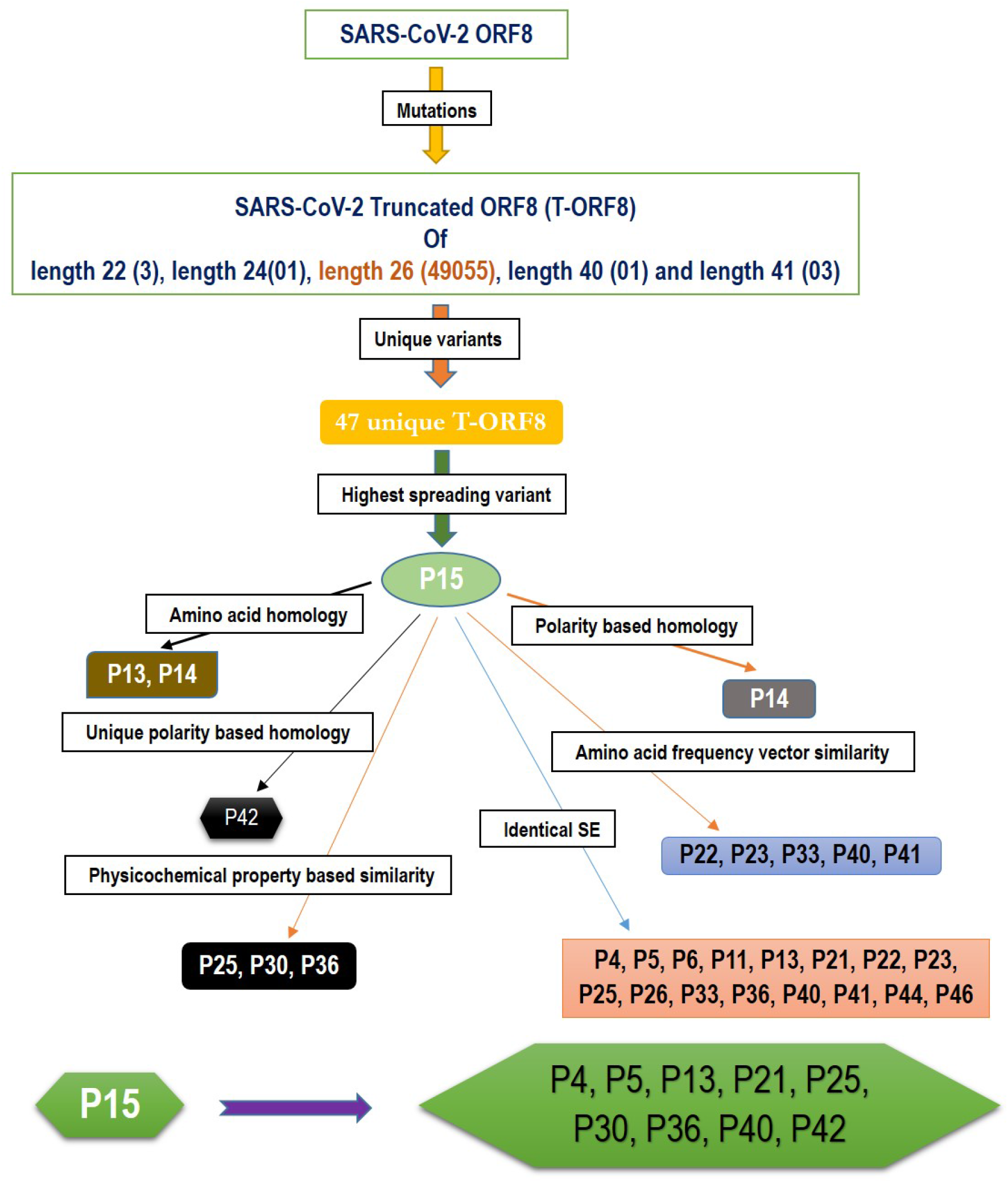
A schematic representation of a possible cluster of unique T-ORF8 variants which were residing in the likelihood of P15 variant. Note: the frequency of each length of T-ORF8 protein was mentioned in parentheses. T-ORF8 variants mentioned in each box were found in the close likelihood of P15 T-ORF8 variants.

All these nine unique T-ORF8 variants had unique polar/non-polar sequences as discussed in Table 7. In addition to the P15 variant, these possible nine emerging variants are likely to appear in the B.1.1.7 lineage of SARS-CoV-2 in near future. As of May 22nd, 2021, it was observed that 16 of 17 COVID-19 affected patients from India (mostly from Gujrat), were infected by the B.1.1.7 lineage of SARS-CoV-2 with the P15 variant, and only one patient (Accession: QVO43928) infected on February 28, 2021 with SARS-CoV-2 strain with the P34 T-ORF8 variant, which had an identical polar/non-polar sequence as that of P15.

## 4. Discussion and Concluding Remarks

ORF8 is 121-amino-acid with two genotypes (orf8L and orf8S), Ig-Like fold, highly immunogenic, SARS-CoV-2 protein interacting with 47 human proteins 15 of them are drug targeting was noticed to interact with MHC-I molecules and significantly down-regulate their surface expression on various cell types [16, 49, 50]. As a result, it was proposed that inhibiting ORF8 function could boost special immune surveillance and speed up SARS-CoV-2 eradication in vivo [50]. ORF8 is not a viroporin like ORF3a of both SARS-CoV-1 and SARS-CoV-2 which are ion channels (viroporins) implicated in virion assembly and membrane budding. So, the viruses lacking E and ORDF3a are not viable and full-length E and ORF3a proteins are required for maximal SARS-CoV replication and virulence [51, 52, 53]. It seems that the ORF8 has only a minor or non-impact on these activities and/or SARS-CoV-2 life cycle as it can survive without functional ORF8, due to many mutations and truncations raised in its gene and protein as above mentioned [23, 54]. The Q27STOP mutation in the ORF8 protein has been discovered to cause 47 distinct truncated ORF8 variations. Furthermore, other truncated protein variants of different lengths 22, 24, 40, and 41 amino acids, were detected, although the frequency of occurrences of those variants was significantly low (Table 2). In Colorado, one T-ORF8 variant of length of 24 amino acids was noticed very recently on April 24, 2021, and this variant is likely to spread further in the future. Other truncated ORF8 variants of amino acids lengths of 22, 40, and 41 no longer appeared in new strains of SARS-CoV-2.

Quantitative characteristics of the 47 unique truncated ORF8 protein variants were examined. All 47 T-ORF8 variants were found in North America, and only the P15 T-ORF8 variant was spread over four continents: Africa, Asia, Europe and South America, until May 22, 2021. In this regard, it is pertinent to raise the question of whether there is any correlation between the spreading of all unique T-ORF8 variants and the epidemiological nature of North America. Within North America, Wang et al. reported that one of the top mutations, 27964C>T-(S24L) on ORF8, has an unusually strong gender dependence [54]. The spread of the P15 variants over the 57 geo-locations across North America was noticed, and in addition, many patients from Asia, Africa, South America and Europe were infected by the particular B.1.1.7 variant of SARS-CoV-2 which contains the P15 variant. Like in many states of the US, also in the US territory of Guam and in North Dakota, most of the patients were noticed to have contracted the P15 variant of the B.1.1.7 lineage. Consequently, the present trend implies that a much higher spread of this lineage with this particular P15 variant is likely to occur. After Europe, Maryland was the first US state to notice the first B.1.1.7 variant with the T-ORF8 P15, but although later this strain remained limited in Maryland it spread further over to other states, such as Florida and Minnesota (Table 5). Furthermore, this analysis reports a set of nine most likely T-ORF8 variants P4, P5, P13, P21, P25, P30, P36, P40, and P42, which were found to be residing in close vicinity of the P15 ORF8 variant. It was noticed that among 47 unique T-ORF8 variants, 28 of them had identical polar/non-polar sequences to that of the P15 variant. Considering the ability of the P15 variant to spread one can assume that the 28 variants with identical polar/non-polar sequence may spread in the near future and cause third, fourth, and fifth waves of COVID-19. As evidence, one patient from India was infected with SARS-CoV-2 with the P34 variant, which has the same polar/non-polar sequences as the P15 variant, as of May 22, 2021 (NCBI accession: QVO43928). The fact that T-ORF8 is still operating as ORF8, is an open issue that needs to be addressed. Reports try to link these T-ORF8 present in many lineages and to COVID-19 patient severity and/or outcomes, effects that contribute to disease progression if associated with the mutations in spike protein [18, 55, 56]. It also is reported that patients infected with SARS-CoV-2, lacking the majority of ORF8 (382 bases), have a lower risk of aggravation, a conclusion supported by Esper et al that accrued variants in spike,ORF8, and ORF3a which were associated with improved clinical outcomes [57]. More recently, SARS-CoV-2 strains were isolated from Washington state, with a stop mutation generating a novel truncated and much smaller ORF8 protein, as well as Hong Kong, which completely missed ORF8 (gene, protein and antibody) and ORF7a, ORF7b [58, 59, 60]. However, the in vitro analysis on Nasal Epithelial cells (NECs) infected with one of these isolates (ORF8 Δ382) may reverse this conclusion as there are no functional significant differences between wild ORF8 and Δ382 [50]. In contrast, Vero-6 cell inoculated with same strain (ORF8 Δ382), showed significantly higher replicative fitness in vitro than the wild type, while no difference was observed in patient viral load, indicating that the deletion variant viruses retained their replicative fitness [61]. In any case, the combinatorial clinical effects of T-ORF8 need to be investigated and analyzed in depth. It is necessary to investigate in detail the functions of T-ORF8 on inflammation and antigen-presenting ability. Finally, caution should be paid for ORF8 as a diagnostic marker as many immunoassay tests depend on its antibody and in light of our analysis for T-ORF8 distribution over different continents [62]. A systematic analysis of its peptide map to determine the effects of these mutations/truncations on the diagnostic potential of the ant-ORF8 antibodies.

## Acknowledgement

SSH devised the study. SSH, VNU, and VK contributed to the implementation of the research, to the analysis of the results. SSH wrote the initial draft of the manuscript. SSH, EMR, KL, PPC, TM, KT, RK, AL, ASA, GKA, AAAA, GP, GC, PA, and MT reviewed and edited. AMB, DB, WBC provided constructive reviews and suggestions. All authors read final version and approve.

## Notes

### Competing Interest Statement

The authors have declared no competing interest.

### Summary of Updates

One new result with Figure 5 has been included.

